# The SARS-CoV-2 nucleocapsid protein preferentially binds long and structured RNAs

**DOI:** 10.1101/2021.12.25.474155

**Authors:** Christen E. Tai, Einav Tayeb-Fligelman, Sarah Griner, Lukasz Salwinski, Jeannette T. Bowler, Romany Abskharon, Xinyi Cheng, Paul M. Seidler, Yi Xiao Jiang, David S. Eisenberg, Feng Guo

**Affiliations:** Department of Biological Chemistry, David Geffen School of Medicine, UCLA, Los Angeles, CA 90095, USA; Department of Chemistry and Biochemistry, UCLA, Los Angeles, CA 90095, USA; Howard Hughes Medical Institute, UCLA, Los Angeles, CA 90095, USA; UCLA-DOE Institute, UCLA, Los Angeles, CA 90095, USA; Molecular Biology Institute, UCLA, Los Angeles, CA 90095, USA

**Keywords:** coronavirus, phosphoprotein, intrinsically disordered region, protein-RNA interaction, RNA-binding protein

## Abstract

The SARS-CoV-2 nucleocapsid protein (NCAP) functions in viral RNA genome packaging, virion assembly, RNA synthesis and translation, and regulation of host immune response. RNA-binding is central to these processes. Little is known how NCAP selects its binding partners in the myriad of host and viral RNAs. To address this fundamental question, we employed electrophoresis mobility shift and competition assays to compare NCAP binding to RNAs that are of SARS-CoV-2 vs. non-SARS-CoV-2, long vs. short, and structured vs. unstructured. We found that although NCAP can bind all RNAs tested, it primarily binds structured RNAs, and their association suppresses strong interaction with single-stranded RNAs. NCAP prefers long RNAs, especially those containing multiple structures separated by single-stranded linkers that presumably offer conformational flexibility. Additionally, all three major regions of NCAP bind RNA, including the low complexity domain and dimerization domain that promote formation of NCAP oligomers, amyloid fibrils and liquid-liquid phase separation. Combining these observations, we propose that NCAP-NCAP interactions that mediate higher-order structures during packaging also drive recognition of the genomic RNA and call this mechanism recognition-by-packaging. This study provides a biochemical basis for understanding the complex NCAP-RNA interactions in the viral life cycle and a broad range of similar biological processes.

**HIGHLIGHTS:**

- NCAP primarily binds structured RNAs.
- NCAP prefers multiple RNA structures separated by single-stranded linkers.
- NCAP favors binding to long RNAs.

## INTRODUCTION

The SARS-CoV-2 virus is a single-stranded (ss), positive-sense virus with an RNA genome of about 30 kb (1, 2). Upon entering host cells, the virus copies its genome by generating the full-length negative-sense genomic RNA (gRNA) for the synthesis of the positive-sense gRNA (3). The positive-sense gRNA also undergoes discontinuous synthesis of negative-sense subgenomic RNAs, which are used to transcribe positive-sense subgenomic messenger RNAs (sgmRNAs) for production of viral structural proteins (3–6). Discontinuous synthesis of subgenomic RNAs is mediated by a transcription regulatory sequence (TRS) that facilitates the fusion of the 5’-leader to each gene. For instance, the NCAP sgmRNA (NsgmRNA) encodes the nucleocapsid protein (NCAP) and contains the 5’ leader sequence and poly(A) tail common to all sgmRNAs. NsgmRNA is the most abundant sgmRNAs in the SARS-CoV-2 transcriptome (7). To produce replication-competent viral particles, it is critical for viral protein(s) to distinguish gRNA from sgmRNAs and host RNAs. The selection of gRNA is thought to be achieved by NCAP and/or the membrane glycoprotein (another structural protein). Although putative packaging signals in the SARS-CoV-2 gRNA have been suggested based on other coronaviruses (8–12), none has been experimentally confirmed.

NCAP is an abundantly expressed and conserved structural protein involved in viral genome packaging and virion assembly (13, 14). Coronavirus NCAPs are known to dimerize and form higher-order oligomers and assemblies (15–18). These properties have been confirmed for the SARS-CoV-2 NCAP, with the dimerization equilibrium constant estimated to be around 0.4 μM (19–21). Consistent with this estimate, a monomeric species has been confirmed at low (100 pM) concentration (22). NCAP can be roughly divided into three major regions: an RNA-binding domain (RBD), a low complexity domain (LCD, an intrinsically disordered linker), and a dimerization domain (DD) (Figure 1A) (17, 19, 20, 22). The RBD and DD are flanked by two additional intrinsically disordered regions (IDRs) called the N-terminal tail (NTT) and C-terminal tail (CTT), respectively. Three-dimensional structures of both the RBD and DD have been determined (23, 24). The latter revealed an extensive dimerization interface, suggesting that NCAP forms a strong dimer. The CTT can help NCAP to further assemble into a tetramer and higher oligomer (25). Both RBD and DD have been shown to bind RNA (22). Upon binding RNA and assembling into virion, NCAP forms higher-order structures as revealed by cryo electron tomography studies (18, 26).

**Figure 1.**
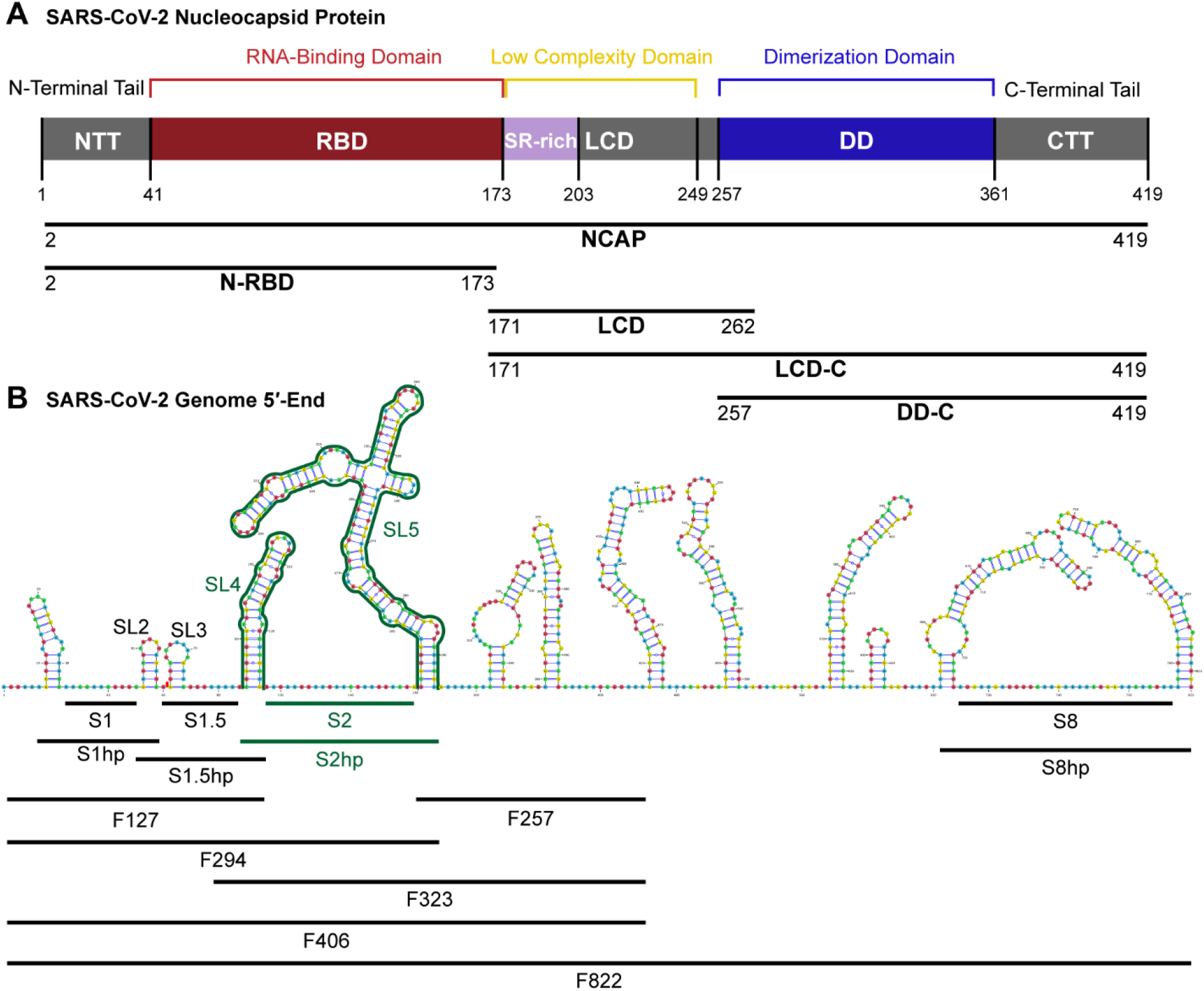
SARS-CoV-2 nucleocapsid protein and RNA constructs used. (**A**) NCAP can be roughly divided into three main regions, the N-terminus-containing RNA-binding domain (N-RBD), the C-terminus-containing dimerization domain (DD-C), and a low complexity domain (LCD) linking the two outer regions. The protein constructs used in this study are shown as bars under the domain diagram. (**B**) The 5’ 822 nt fragment of the SARS-CoV-2 genome (F822) contains multiple structured regions linked by single-stranded RNA. The RNAs used from this region included single-stranded Site 1, 1.5, 2, and 8 as well as RNA constructs containing the flanking hairpin regions (S1hp, S1.5hp, S2hp, S8hp). Fragments of the 5’ 406 nt (F406) including F127, F257, F294, and F323 are also shown. The primary focus of this study is on S2hp and its truncations, including stem loop 4 and 5 (SL4, SL5) and S2 (highlighted in green). The RNA secondary structure was previously established with inline and RNase V1 probing, NMR spectroscopy, and SHAPE-MaP (27, 34, 57) and was visualized using VARNA (58).

The NCAP-RNA interaction is complex. Previous research using biolayer interferometry has shown that a truncation of NCAP containing the DD-C region can bind short, ss and double-stranded (ds) RNAs containing the TRS with *K*_D_s in the 6–33 μM range (20). Iserman *et al.* identified many NCAP-crosslinking sites within the 5’-end 1000 nt of the SARS-CoV-2 genome (27, 28). The most pronounced NCAP-crosslinking sites were listed as Sites 1–10 (S1–S10), which include mostly ss regions and some A/U-rich structured regions. When the molar ratio of NCAP and the 5’-end RNA was kept at 20:1, the ss Sites 2 and 8 served as the principal NCAP-binding sites. As the molar ratio increased to 80:1 and 160:1, other sites were occupied. This increase in valency was accompanied by the transition of the complexes from diffuse to condensed states. The authors suggested that the stem loop structures flanking the ss NCAP-binding sites may be important for the protein-RNA interaction.

Upon binding RNA, NCAP can form liquid-liquid phase separation (LLPS), which is thought to locally concentrate viral components to facilitate viral genome packaging and to exclude many cellular proteins as a means of protection from the host immune system (29, 30). The 5’-end region of SARS-CoV-2 genomic RNA has been shown to stimulate LLPS and increasing RNA length promotes N-protein to LLPS (27). In a parallel study, we show that NCAP can form amyloid fibrils (31). A recent preprint showed that the dsRNA regions flanking the principal NCAP-binding sites drive the formation of LLPS with NCAP (32). However, because these studies emphasized how NCAP and RNA form LLPS, it remains unknown how relevant RNA features and elements contribute to NCAP-binding affinity and specificity.

Here, we characterized NCAP-RNA interactions in-depth using biochemical methods. SARS-CoV-2 and non-SARS-CoV-2 RNAs of differing lengths and structures were compared in affinity measurements and competition assays (Figure 1B). These RNAs included both ss sites with and without flanking hairpins (stem loops), with a focus on the most prominent NCAP-crosslinking Site 2 (S2) and the flanking stem loops 4 and 5 (SL4, SL5). The latter RNA, which we call S2hp (Site 2 with flanking hairpins), is also interesting because it contains the translation start site of ORF1a and two upstream open reading frames (uORFs) that may be functionally important (33, 34). Surprisingly, we found that NCAP primarily binds the structures surrounding the ss crosslinking sites. In addition, we demonstrate a general preference of NCAP for long RNAs and propose a mechanism to explain how NCAP favors the long viral gRNA.

## MATERIALS AND METHODS

### Molecular biology reagents

Phusion HF DNA polymerase, restriction enzymes, Quick CIP, and T4 Polynucleotide Kinase (PNK) were obtained from New England BioLabs. Custom DNA oligonucleotides and gBlocks were synthesized by Integrated DNA Technologies. RNA oligos were synthesized by Horizon. Acrylamide:Bis-Acrylamide (29:1 ratio) 40% stock solution used in electrophoresis was purchased from Fisher Scientific.

### *In vitro* transcription and purification

The sequences of S1hp, S1.5hp, and S2hp were cloned from a gBlock containing the 5’ 1,000 nt of SARS-CoV-2 genome into the pUC19 vector using restriction sites EcoRI and KpnI. Two rounds of PCR were used to amplify and merge the 5’-leader (from the gBlock) and the NCAP-coding sequence (from 2019-nCoV Control Plasmid (IDT Inc, Cat No 10006625)), to produce the NsgmRNA sequence, which was subsequently cloned into the pUC19 vector using a single restriction site KpnI. The NsgmRNA used in this study does not include the 6 nt at 5’ end, 3’-UTR, and poly(A) tail. The sequences of all cloned constructs were confirmed using Sanger sequencing.

Other than synthetic oligos, all RNAs were produced via *in vitro* transcription. The DNA templates were prepared using PCR with the Phusion HF DNA polymerase. The PCR templates were either cloned plasmids or gBlock DNAs. The forward primers used in these PCR reactions contained a biotin at their 5’-end so that the DNA template can be removed after transcription using streptavidin beads. PCR products were purified via anion exchange chromatography using a 5-mL HiTrap Q HP column (Cytiva). The running buffers contained 10 mM Na HEPES pH 7.0 and either 10 mM (buffer A) or 2 M (buffer B) NaCl. Purified PCR templates were concentrated using Amicon Ultra-4 centrifugal filter units (MilliporeSigma) and buffer-exchanged to 10 mM Na HEPES pH 7.0.

Transcription reactions ranging from 5 to 40 mL in volume were set up and incubated at 37°C for one hour. Streptavidin agarose beads (ThermoFisher) were then added to the reactions, which were mixed on a rotator at room temperature for 10 min. Subsequently, the solutions were centrifuged at 100xg for 10 min at 4°C. The supernatant was immediately loaded onto HiTrap Q HP columns, washed, and purified. The purified RNA was concentrated using Amicon Ultra-15 centrifugal filter units (MilliporeSigma) and buffer-exchanged to 10 mM Na HEPES pH 7.0. The purity of the RNAs was confirmed using denaturing polyacrylamide gels.

### Protein expression and purification

Detailed protein expression and purification procedures are described in another manuscript (31). Briefly, the coding sequences for NCAP and its truncations with N-terminal 6xHis-SUMO tag were cloned into the pEC-28a vector. The proteins were expressed in *E. coli* Rosetta2 (DE3) cells and were first purified using Ni affinity chromatography. The His-SUMO tag was cleaved off and the cleaved proteins were further purified using reverse-His tag purification and size exclusion chromatography. SDS-PAGE and 260/280 nm absorbance ratio were used confirm protein purity and low RNA contamination, respectively. The purified proteins were concentrated in 20 mM Tris pH 8.0 and 300 mM NaCl, and stored at −80°C.

### Electrophoresis mobility shift assay (EMSA)

Interactions between NCAP constructs and RNAs were analyzed using 5–8% native polyacrylamide gels run at room temperature in 1xTBE buffer. S2hp, SL4, and SL5 generated via *in vitro* transcription were treated with Quick Calf Intestinal Alkaline Phosphatase before labeling with [γ-^32^P]-ATP (Perkin Elmer) and T4 PNK. The synthetic S2 was directly labeled with [γ-32P]-ATP and T4 PNK. The labeled RNAs were purified using Microspin G-25 columns (Cytiva). Each 10-μL binding reaction contained 1xPBS buffer, trace amounts (<0.5 nM, true for all assays in this study) radiolabeled RNA, various concentrations of NCAP proteins, with tRNAs and competitor RNAs as indicated in figures and tables. All binding reactions were mixed well and incubated at room temperature for at least an hour before mixed with 6x native RNA loading dye (15% Ficoll, 0.02% xylene cyanol, 1xPBS) and loaded onto custom-poured mini gels (BIO-RAD). All gels were run at room temperature, dried, and exposed to a phosphorimager screen. The gel images were obtained using an Amersham™ Typhoon™ scanner (Cytiva) and were quantified using Quantity One (BIO-RAD, version 4.6.9) or ImageJ. The data were fitted and plotted using Prism (GraphPad, version 8). For *K*_D_ measurements, Y = Bmax * X^n^ / (*K*_D_^n^ + X^n^), where Y is the fraction of RNA bound to NCAP and X is the NCAP concentration. n is the Hill coefficient for which the values of 1,2, 3, and 4 were tried and the value resulting in the best fit to data was used.

### Competition assay

Master mixes containing radiolabeled RNA, 4 μM NCAP, and 1xPBS were made and distributed into individual reactions. 4 μM tRNAs were also added for all competition assays with radiolabeled S2hp, whereas no tRNAs were used for radiolabeled S2. Serial dilutions of each unlabeled competitor RNA were added, and samples were incubated at room temperature for at least an hour before loading onto gels. Gels were run, dried, and processed as stated in EMSA.

### Protein saturation assay

Master mixes of radiolabeled RNA with the same unlabeled RNA were made in 1xPBS. The unlabeled RNA concentration chosen were well above the measured *K*_D_ values. Serial dilutions of each protein were added, and samples incubated at room temperature for at least an hour before loading onto gels.

### Statistical analyses

Unpaired Student’s *t*-tests with two tails were performed in Excel (Microsoft).

## RESULTS

### SL5 makes a major contribution to the binding affinity of S2hp for NCAP

We characterized the interaction between NCAP and viral RNA fragments using electrophoresis mobility shift assays (EMSA). This assay was previously used to study the interaction between SARS-CoV-1 NCAP and a 20-nt poly(U) ssRNA (35). We initially focused on RNA fragments centered on the previously reported prominent NCAP-binding S2, which consists of an A/U-rich, 22 nt ss region. Fragments studied also include S2hp, which contains S2 and flanking stem loop regions, as well as the individual stem loops, SL4 and SL5 (Figure 1B). EMSA was used to determine affinities (reported as the inversely related *K*_D_ values) by mixing trace amounts (< 0.5 nM) of radiolabeled RNA with increasing concentrations of NCAP. The gels showed that the free RNA bands disappeared as the NCAP concentration was increased, indicative of RNA-NCAP complex formation (Supplementary Figure S1). Some smeared intensities were observed at the binding transition, likely resulting from transition complexes. The majority of the RNA-NCAP complexes were stuck in the wells, preventing reliable detection, similarly observed previously for the SARS-CoV-1 NCAP (35). Thus, the *K*D was calculated using the disappearance of free RNA as a proxy for the formation of RNA-NCAP complexes. Whereas the data did not fit in a 1:1 binding model, cooperative models with Hill coefficients of 2 to 4 fit the data well, giving *K*_D_ values summarized in Table 1. The affinities of NCAP for S2hp (*K*_D_ = 173 ± 14 nM, mean ± SD) is significantly higher than that for S2 (*K*_D_ = 216 ± 23 nM), with *P* value of 0.02, indicating that the flanking stem loop structures are important for NCAP binding. Indeed, the affinity for SL5 (*K*_D_ = 166 ± 36 nM) is similar to that for S2hp, whereas SL4 binds NCAP the weakest, with *K*_D_ of 356 ± 43 nM. The free SL4 RNA migrated as two bands, likely corresponding to hairpin and dimeric duplex conformations, despite different annealing procedures tried (Supplementary Figure S1C). We obtained *K*_D_ values by either combining the intensities of the hairpin and duplex bands as that of free RNA or keeping them separate (Supplementary Figure S1D). The affinity for the hairpin is about half that for the duplex, and they bracket that obtained from the combined intensities (Table 1). Fortunately, in this study regardless how SL4 is quantified, our conclusions are unchanged. In addition, although SL4 has a NCAP affinity significantly different from those of S2hp and its other truncations, the difference is in about three-fold range (*K*_D_’s of 166–569 nM). These moderate differences nevertheless suggest that SL5 is a major contributor to NCAP-binding affinity of the S2hp region.

**Table 1.** Summary of binding affinities of NCAP for viral RNA fragments.

| *RNA* | *Length (nt)* | *K<sub>D</sub> (- tRNAs)* | *N* | *K<sub>D</sub> (+ tRNAs)\** | *N* | *K<sub>D</sub>(+ tRNAs) / K<sub>D</sub>(- tRNAs)* |
| --- | --- | --- | --- | --- | --- | --- |
| S2 | 22 | 216 ± 23 nM | 5 | *n.d.* |  | *n.d.* |
| SL4** | 44 |  | 3 |  | 3 |  |
| Combined |  | 356 ± 43 nM |  | 4.8 ± 0.2 μM |  | 13 |
| Hairpin |  | 569 ± 67 nM |  | 8.6 ± 1.0 μM |  | 15 |
| Duplex |  | 235 ± 27 nM |  | 3.9 ± 0.5 μM |  | 17 |
| SL5 | 145 | 166 ± 36 nM | 7 | 1.3 ± 0.1 μM | 4 | 7.5 |
| S2hp | 211 | 173 ± 14 nM | 4 | 312 ± 39 nM | 4 | 1.8 |
| F406# | 408 | 1.2 ± 0.2 nM | 3 |  |  |  |
| F323# | 325 | 29 ± 9 nM | 3 |  |  |  |
| F294# | 296 | 26 ± 10 nM | 3 |  |  |  |
| F257# | 259 | 23 ± 5 nM | 3 |  |  |  |
| F127# | 129 | 183 ± 56 nM | 3 |  |  |  |

N refers to the number of repeats performed.
\* Measured in the presence of 4 μM tRNAs. The presence of tRNAs abolished binding of S2 to NCAP so *K<sub>D</sub>* could not be determined. *K<sub>D</sub>* values for F series of RNAs were measured in the absence of tRNAs.
\*\* Free SL4 RNA migrates as two bands, which I believe respond to hairpin and dimeric duplex conformations. The bands quantified either combined or separately.

### RNA structure and length determine competitive binding to NCAP

One limitation of the EMSA was the poor migration of RNA-NCAP complexes into gels. To address this issue, we added varying concentrations of yeast tRNAs to the binding reactions. We found that yeast tRNAs helped the S2hp-NCAP complex migrate into the gel as a single band, with 1 to 4 μM needed to achieve maximal effects (Figure 2A and B). A control reaction indicated that yeast tRNAs do not alter the mobility of S2hp in the absence of NCAP (Figure 2C, lanes 13 and 14). It is likely that yeast tRNAs directly interact with NCAP and prevent its aggregation. Importantly, tRNAs are highly abundant in the cytoplasm where SARS-CoV-2 replicates and NCAP interacts with viral RNAs. Therefore, besides better visualization of RNA-NCAP complexes, binding assays in the presence of tRNAs reflect interactions closer to physiological conditions. Since yeast and human tRNAs share similar folds (36), we added 4 μM of yeast tRNAs (simply called tRNAs from now on) to binding reactions in subsequent experiments where noted. As later results (Figure 5E, lanes 4, 6, 8) show, other RNAs can also help S2hp-NCAP complex migration into gels, so the effect on NCAP-RNA gel migration is not specific to tRNAs.

**Figure 2.**
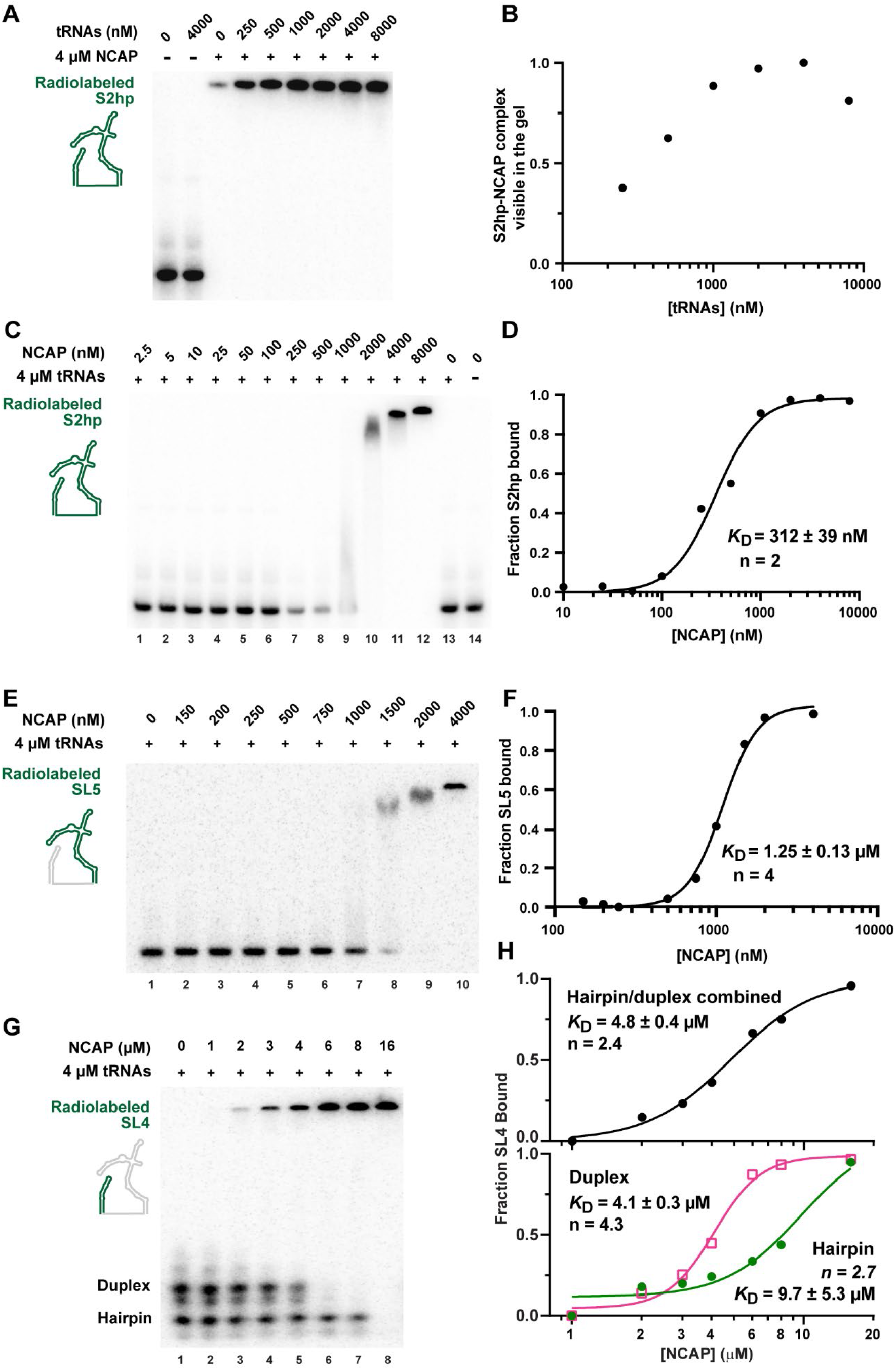
Binding affinities of NCAP for S2hp fragments in the presence of tRNAs. (**A**) EMSA of radiolabeled S2hp with 4 μM NCAP and varying concentrations of unlabeled tRNA competitors. tRNAs increase RNA-NCAP complex migration into gels. (**B**) Quantification shows that 1–4 μM tRNAs achieve maximum signals from the radiolabeled S2hp in complex with NCAP. (**C**) EMSA of radiolabeled S2hp with 4 μM tRNAs and varying concentrations of NCAP. These conditions were kept throughout measuring *K*_D_ of S2hp truncations in the presence of tRNAs. (**D**) S2hp binds NCAP with a *K*_D_ = 312 ± 39 nM. n represents Hill coefficients used for fitting. (**E**) EMSA of radiolabeled SL5. (**F**) Quantification and fitting show that SL5 binds NCAP with a *K*_D_ = 1.3 ± 0.1 μM. (**G**) EMSA of radiolabeled SL4. (**H**) Quantification of the gel shown in (G). The free SL4 bands were quantified either combined (top panel) or separately as duplex and hairpin (bottom panel). In curve fitting, the Hill coefficient was allowed to vary during the fitting. The experiments were repeated 3–4 times and the results are summarized in Table 1. Shown here are representative gel images and quantification results. Curving fitting provides the *K*_D_ values with standard errors.

We measured the binding affinities of NCAP for S2hp, SL5 and SL4 in the presence of tRNAs and observed pronounced contributions by RNA structure and length. The presence of tRNAs allows all three complexes to migrate into gels and significantly decreased the apparent affinities (Figure 2C–H and Table 1). The *K*_D_ values are 312 ± 39 nM for S2hp, 1.25 ± 0.13 μM for SL5, and 4.8 ± 0.2 μM for SL4 (with two free RNA bands combined). The affinity for SL4 hairpin is again about half that for the duplex (Figure 2H and Table 1). For S2, no binding was detected with 4 μM NCAP in the presence of tRNAs (data not shown). Whereas the apparent affinity for S2hp is highest, that for S2 is the lowest, corroborating the idea that the structured regions neighboring S2 are important for binding NCAP. We note that in the presence of tRNAs, the apparent affinity follows the order of the RNA length, with S2hp the highest. The fold affinity decreases caused by tRNAs also follow the trend of RNA length, with 1.8-fold for the 213-nt S2hp, 7.5-fold for the 147-nt SL5, 13–17-fold for the 46-nt SL4, and a greater (despite undetermined) value for the 22-nt S2. These results provide the initial indication that structure and length define RNA-binding preference of NCAP.

Operationally, competition assays are highly suitable for investigating RNA-binding specificity of proteins. Thus, we compared competitive binding of different viral RNA fragments to NCAP. In the first set of assays, a trace amount of ^32^P-labeled S2hp was mixed with NCAP, tRNAs, and RNA competitors, including single-stranded NCAP-crosslinking sites revealed by the previous study (S2, S1, S1.5, and S8) (27) and their expanded constructs containing flanking hairpins (S2hp, S1hp, S1.5hp, and S8hp) (Figure 1B). We added S1.5 because it contains the TRS sequence. S1.5 contains SL3, which only contains four base pairs in the stem and is likely to be marginally stable. tRNAs were added to allow the S2hp-NCAP complex to migrate into gels. As expected, unlabeled S2hp was able to compete with the labeled S2hp, disrupting the complex and returning the labeled RNA mostly to the free state (Figure 3A, lane 4). In contrast, 4 μM single-stranded S2, S1, S1.5, or S8 failed to disrupt the complex (Figure 3A, lanes 5, 7, 9, 11). Among the other hairpin-containing competitors, only S8hp was able to disrupt the S2hp-NCAP complex, creating two smeary bands with greater mobility (Figure 3A, lane 10). No free S2hp band was observed, indicating that the NCAP protein is still able to bind S2hp to some extent. S8hp (172 nt) contains two large (> 44 nt) stem loop structures flanking a single-stranded region and is closest in size to S2hp (211 nt), highlighting the importance flanking structures and length in binding NCAP. These results recapitulate the previous finding by Iserman et al. that S2 and S8 are the most strongly crosslinked by NCAP (27).

**Figure 3.**
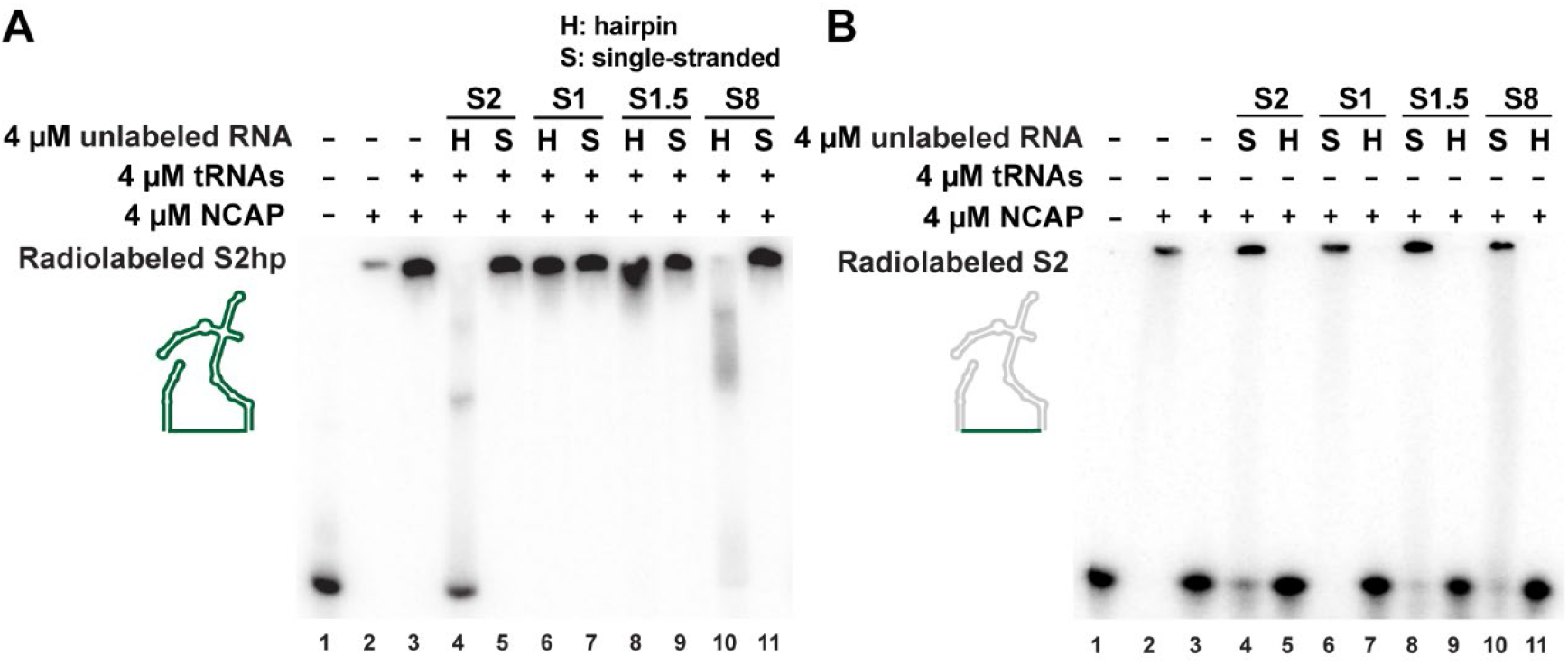
S2hp and S2 in competition with other 5’ SARS-CoV-2 RNAs. (**A**) S2hp outcompetes other 5’ viral RNAs with single-stranded and hairpin structures. (**B**) S2 outcompetes single-stranded viral RNAs, including the longer S1.5 and S8 constructs. S2 is outcompeted for binding to NCAP by longer viral RNAs with flanking hairpin structures.

Our conclusion that NCAP prefers binding of structured RNA over unstructured RNA is supported by parallel competition assays in which ^32^P-labeled S2 was used instead of S2hp, in the absence of tRNAs. All RNAs containing flanking hairpin regions outcompete S2 for binding to NCAP as evidenced by the free S2 RNA bands (Figure 3B, lanes 3, 5, 7, 9, 11). At the same molar concentration (4 μM), none of the single strands, including S2 itself, can efficiently disrupt the radiolabeled S2-NCAP complex (Figure 3B, lanes 4, 6, 8, 10).

### Flanking structures of S2hp harbor the primary NCAP-binding sites

To understand the interactions responsible for the RNA-binding selectivity of NCAP, we determined the NCAP-binding stoichiometry of S2hp and its components. Binding reactions containing constant amounts of radiolabeled and unlabeled RNAs, 4 μM tRNAs, and varying concentrations of NCAP were analyzed using EMSA (Figure 4). The trace amount of radiolabeled RNA allowed detection and quantification. The concentration of unlabeled RNA chosen was above the *K*_D_ so that most NCAP proteins were bound unless the RNA had been saturated. The saturation assay with S2hp revealed sequential formation of three complex bands as NCAP concentration increased (Figure 4A). Quantification indicated that the fastest-migrating band forms at lower NCAP concentrations with its intensity peaked at 2:1 NCAP protomer:RNA molar ratio (Figure 4B), suggesting that the complex corresponds to a NCAP dimer bound to S2hp. The complex with medium mobility peaked at between 3:1 and 4:1 molar ratio, likely containing S2hp bound to two NCAP dimers. The slowest-migrating band became prominent at 4:1 ratio and was almost the only band at 5:1, consistent with three NCAP dimers bound to S2hp (Figure 4C). This analysis identified three NCAP dimer binding sites on S2hp.

**Figure 4.**
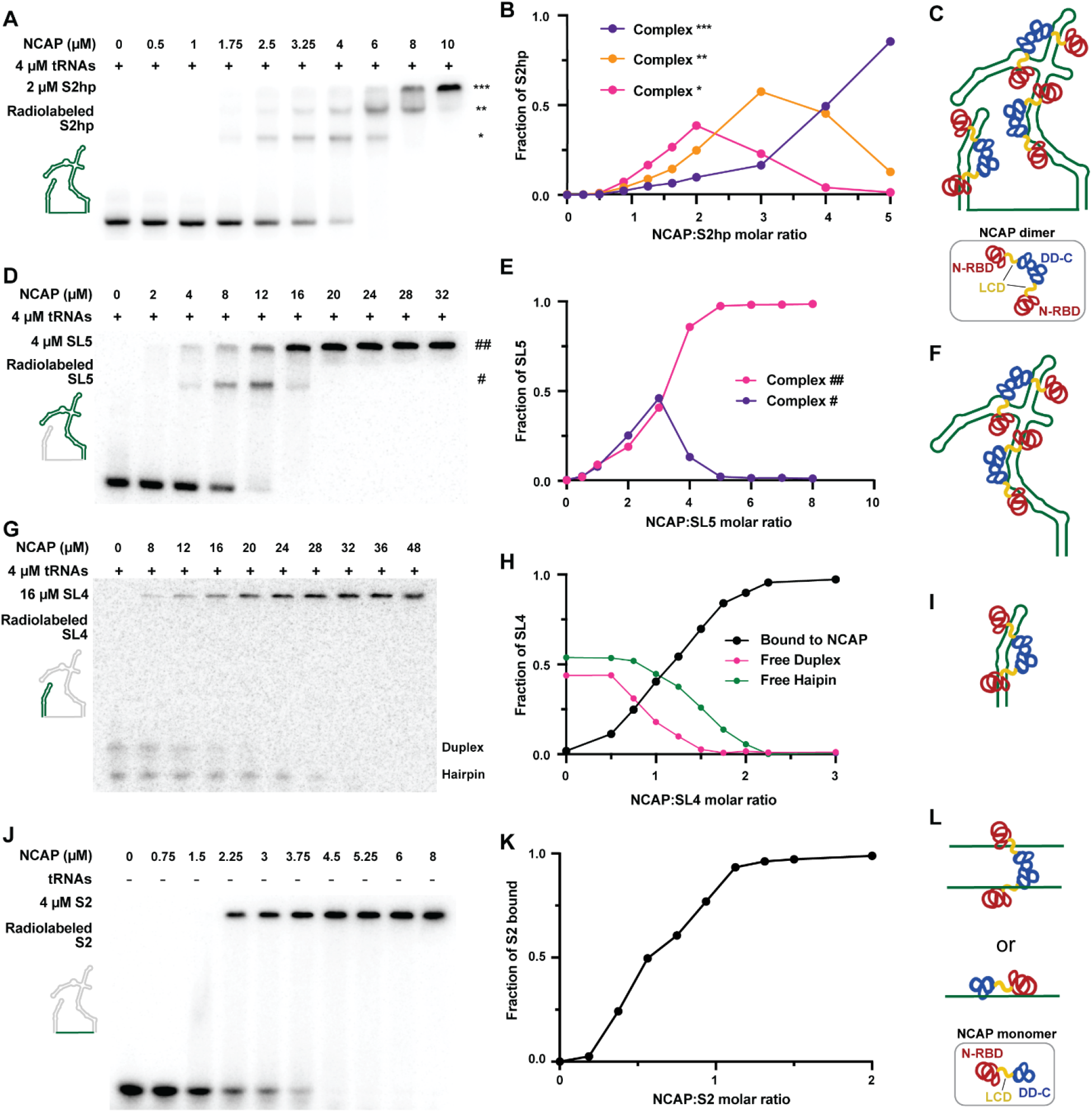
Saturation assays reveal NCAP-binding sites on S2hp. Radiolabeled and unlabeled RNAs, NCAP and tRNAs were mixed at concentrations indicated in the figures. The concentrations of unlabeled RNAs were well above the *K*_D_ values to assure binding. (**A**) Saturation experiment for S2hp shows three distinct NCAP-RNA complexes. (**B**) Quantification of the complexes suggests three NCAP-binding sites on S2hp. A model of the fully assembled S2hp-NCAP complex is shown in (**C**). NCAP is assumed to be a dimer, as illustrated in the box. (**D**) Saturation assay for SL5 shows two NCAP-RNA complexes. (**E**) Quantification of the complexes indicates saturation at 4:1 NCAP protomer:SL5 molar ratio, suggesting that each SL5 molecule can bind two NCAP dimers (**F**). (**G**) Saturation assay for SL4. SL4 folds into two conformations that correspond to duplex and hairpin, respectively. (**H**) Quantification of the SL4-NCAP complex, as well as free duplex and hairpin, indicates saturation at two NCAP protomers per SL4 molecule. (**I**) We suggest that each SL4 binds a NCAP dimer. (**J**) Saturation assay for S2. (**H**) Quantification of the free S2 band and inference of the fraction bound indicate saturation at one NCAP protomer per S2 molecule. (**L**) We suggest that either each NCAP dimer can bind two S2 molecules or one NCAP monomer binds one S2.

Our subsequent experiments located the three NCAP-binding sites to the flanking structures of S2hp. The saturation assays with SL5 showed two complex bands (Figure 4D and E). The intensity of the faster-migrating band peaked between 2:1 and 3:1 NCAP protomer:RNA molar ratio, suggesting that it corresponds to one NCAP dimer bound to SL5. The slower-migrating band became the dominant species at 4:1 molar ratio, indicative of two NCAP dimers bound to SL5 (Figure 4F). This result suggests that SL5 has two NCAP-binding sites. The SL4 assays showed one complex band that was saturated a 2:1 NCAP:SL4 ratio, suggesting that one NCAP dimer binds to each SL4 molecule (Figure 4G, H and I).

We also determined the S2-NCAP binding stoichiometry. Because S2 does not bind NCAP strongly, these saturation experiments were performed in the absence of tRNAs. Increasing concentrations of NCAP were incubated with 4 μM unlabeled S2 and trace but constant amount of radiolabeled S2. The free S2 band disappeared and thus S2 became fully bound with between 3.75 and 4.5 μM NCAP (Figure 4J and K), corresponding to each NCAP protomer binding to one S2 molecule. We think either each NCAP dimer binds two S2 molecules or each NCAP monomer binds one S2 (Figure 4L). This result suggests that there is a surface on NCAP responsible for binding S2 and likely other ss regions. In the S2hp-NCAP complexes, the S2 region may bind to NCAP subunits associated with a flanking structure or is available for recruiting additional NCAP proteins.

### Structured RNAs suppress strong NCAP-ssRNA interaction

We next characterized the ssRNA-binding surface of NCAP using competition assays. In these assays, varying concentration of an unlabeled competitor RNA was incubated with constant concentrations of NCAP and radiolabeled S2. We first used S2hp, SL5 and SL4 as competitors to dissect the interactions of NCAP with different components of S2hp. Approximately 500 nM of S2hp was sufficient to take over the 4-μM NCAP added and return S2 to the free state (Supplementary Figure S2A), suggesting most of the ssRNA-binding surface of NCAP has been occupied in the full NCAP-S2hp complex. SL5 effectively competed with S2, resulting in free S2 at 1 μM (Supplementary Figure S2B). At this concentration, SL5 is also expected to occupy all 4 μM of NCAP. This result suggests that association of SL5 with an NCAP protein precludes it from binding S2.

SL4 displayed partial competence against S2 binding. SL4 released most S2 RNA to the free state at 4 μM and higher concentrations, but some S2-NCAP complex remained even at the highest SL4 concentration of 32 μM (Supplementary Figure S2C). This latter result suggests that NCAP bound to SL4 has some S2-binding capacity. Interpretation of these data is complicated by the fact that SL4 folds into hairpin and dimeric duplex conformations (as seen in Figures 2G and 4C). The duplex has higher affinity to NCAP than the hairpin (Supplementary Figure S1C and Figure 2G), and therefore is likely to contribute more to the competition. Nevertheless, the results clearly indicate that association of NCAP with SL4 hairpin/duplex greatly reduce S2 binding. When the structured RNA is relatively small, a NCAP protein may be able to bind the structured and ss RNAs simultaneously. In the context of S2hp-NCAP complex, these results suggest that the S2 strand can either bind to the NCAP subunit associated with the SL4 site, associate with a NCAP protein other than the three dimers bound to SL5 and SL4, or do not bind any NCAP protein if none is available.

Using the S2 competition assay, we compared synthetic small interfering RNAs (siRNAs) in both ss and duplex forms. The siDGCR8-1 sense strand contains 21 nt with a 5’ phosphate, which is similar in size to the 22-nt S2. The siDGCR8-1 duplex was produced by annealing the sense and antisense strands, forming 19 base pairs with 2-nt overhangs at the 3’ of each strand (37). The siDGCR8-1 duplex is similar in size to the ~17-bp stem of SL4. The results showed that 16 μM of siDGCR8-1 duplex returned nearly all radiolabeled S2 to the free form, whereas at least 64 μM of siDGCR8-1 sense was required (Supplementary Figure S2D and E). This result reinforces the conclusions that NCAP prefers to bind structured RNAs and that their association opposes strong interaction with ssRNAs.

### Combination of structures and ss regions makes an RNA competitive for binding NCAP

We next compared the competitiveness of viral and non-viral RNAs against S2hp-NCAP binding. In these competition assays, varying concentration of an unlabeled competitor RNA was incubated with constant concentrations of NCAP, tRNAs, and radiolabeled S2hp. We started by using S2hp itself as the competitor. Increasing concentrations of unlabeled S2hp gradually disrupted the fully assembled complex (Figure 5A). At 1 μM, unlabeled S2hp produced complexes containing two and three NCAP dimers. At 2 μM, we observed the complex bound with one NCAP dimer and free S2hp. At 4 μM and higher concentration, most S2hp was free. These results provide a benchmark for comparing with other competitors and suggest a hierarchy among the three NCAP dimers during disassembly from S2hp.

**Figure 5.**
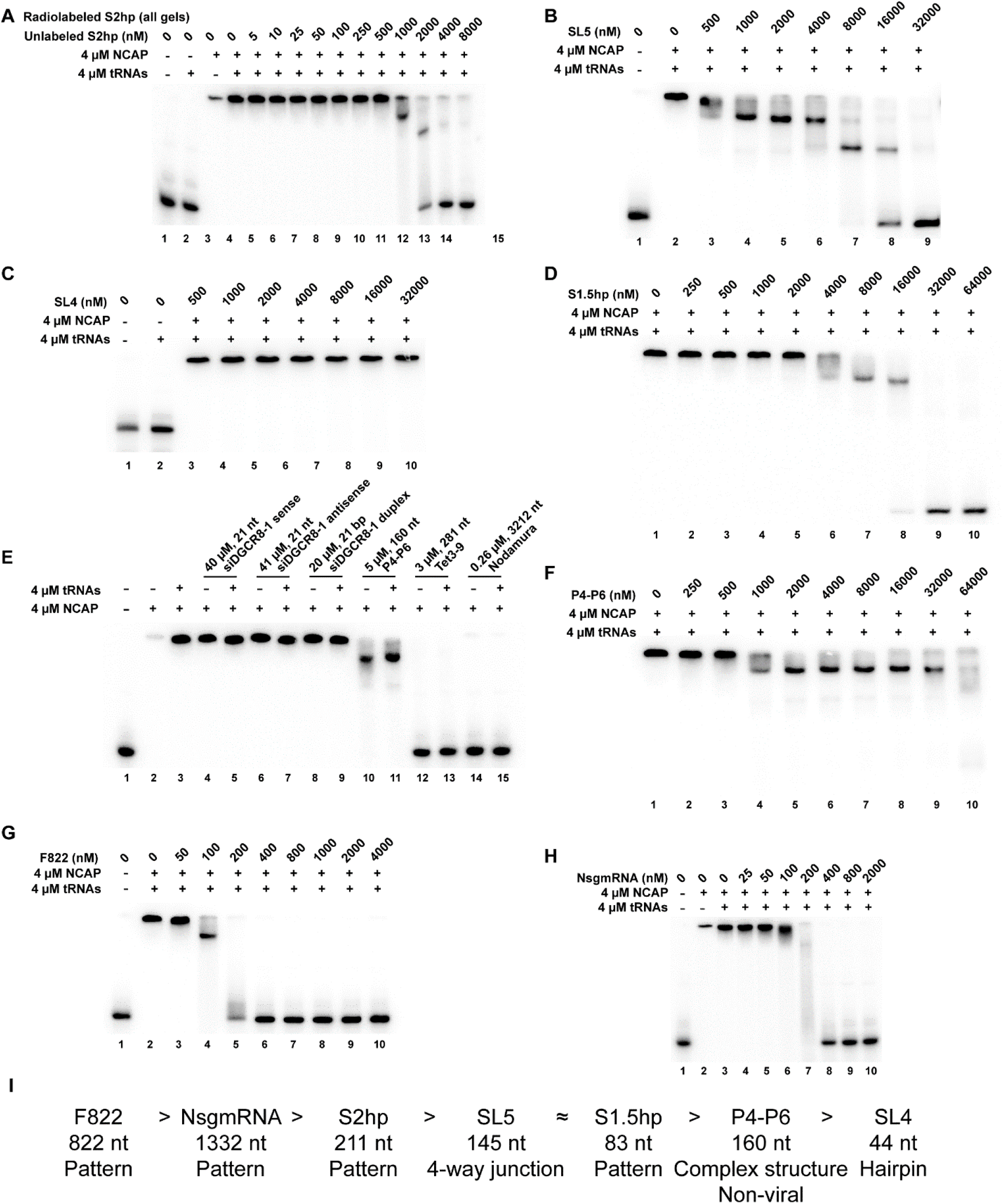
Competition against radiolabeled S2hp binding to NCAP by unlabeled viral and non-viral RNAs of different lengths and structures. Unlabeled S2hp (**A**), SL5 (**B**), SL4 (**C**), S1.5hp (**D**), P4-P6 (**F**), F822 (**G**), and NsgmRNA (1,332 nt) (**H**) were added to the competition reactions. (**E**) Competition gel shift with nonspecific RNA competitors of varying lengths added at equal mass concentration. (**I**) Summary of RNA competitiveness to completely disrupt the S2hp-NCAP complex. The length and structural type are indicated below each RNA. Pattern refers to more than one structures separated by ss linkers.

Subsequent competition assays showed that components of S2hp achieve NCAP-binding competitiveness together. The experiments using unlabeled SL5 also showed hierarchical disassembly of the S2hp-NCAP complex (Figure 5B). At 1–4 μM, SL5 generated mainly the complex containing two NCAP dimers. At 8 μM, the dominating complex contained one NCAP dimer. Roughly equal amounts of free S2hp and complex with one NCAP dimer were produced by 16 μM SL5 (Figure 5B, lane 8), like what 2 μM S2hp accomplished (Figure 5A, lane 13). It took 32 μM SL5 to abolish the complexes (Figure 5B, lane 9), whereas only 4 μM S2hp was needed (Figure 5A, lane 14). This large difference cannot be explained by the lower binding stoichiometry of SL5 and is likely to be caused by its lower affinity for NCAP than that of S2hp. At up to 64 μM, neither SL4 nor S2 can disrupt the S2hp-NCAP complex (Figure 5C and data not shown). These results indicate that individual components of S2hp, namely the SL5, SL4 and S2 regions, are not as competitive as S2hp and thereby must work together to achieve the higher competitiveness for NCAP binding.

Our data support the previous proposal that structures separated by ss regions are an important pattern recognized by NCAP (27). This idea is corroborated by the experiments using S1.5hp as the competitor (Figure 5D). S1.5hp contains SL4 and the upstream SL2, SL3 and the linkers in between (Figure 1B). The linker between SL3 and SL4 contains eight nucleotides. Therefore, S1.5hp fits the structure-ss-structure pattern. In contrast to SL4, S1.5hp reduces the full S2hp-NCAP complex to the one mostly containing two NCAP dimers at 4–16 μM and to free RNA at 32 μM and above. Therefore, addition of structures and a linker to SL4 renders the RNA more competitive in NCAP binding. S1.5hp (84 nt) abolishes S2hp-NCAP interaction at concentrations akin to those of SL5, despite being 63 nt shorter, further highlighting the efficacy of the pattern.

### The length and structural preferences apply to non-SARS-CoV-2 RNAs

To further investigate how length and structure affect RNA binding to NCAP, non-SARS-CoV-2 RNAs of varying lengths and secondary structures were used as competitors against radiolabeled S2hp for binding NCAP (Figure 5E). Each competition reaction was performed with and without tRNAs, with little differences observed. For RNA competitors to block the binding of radiolabeled RNA, their affinity for NCAP as well as their ability and capacity to occupy overlapping sites are critical. To compare more rigorously, we normalized the number of nucleotides by using the same mass concentration (271 ng/μL, equivalent to 4 μM S2hp) for all competitors (Supplementary Table S2). Small RNAs such as single-stranded forms of siDGCR8-1 (sense and antisense, both 21 nt) and siDGCR8-1 duplex (37) cannot outcompete S2hp in binding NCAP (Figure 3C, lanes 4-9). A highly structured RNA P4-P6 (160 nt, a domain of the *Tetrahymena* ribozyme (38)) can partially disrupt the S2hp-NCAP complex, resulting in a faster migrating species rather than free RNA (lanes 10, 11). The mobility of this species suggests that it is a complex of S2hp bound to two NCAP dimers. Two RNAs larger than S2hp, Tet3-9 (281 nt, another highly structured *Tetrahymena* ribozyme construct (39)) and Nodamura replicon (3212 nt (40)), completely disrupts the binding of S2hp to NCAP as evidenced by the free S2hp band (lanes 12-15). These results show that the SARS-CoV-2 NCAP can bind RNAs not from its own genome and that it has a general preference for longer RNAs.

Although NCAP prefers longer, structured RNAs, not all structures are preferred. The highly structured P4-P6 (160 nt) is between SL5 (145 nt) and S2hp (211 nt) in length. Addition of 1–32 μM competitor P4-P6 can disrupt NCAP binding to S2hp, forming a faster migrating band (Figure 5F, lanes 4-9). However, even at 64 μM, P4-P6 is unable to fully disrupt S2hp binding to NCAP (Figure 5F, lane 10). This result shows that P4-P6 is able to compete with one NCAP-binding site (possibly the weaker SL4 site) but not efficiently with the remaining two in S2hp.

We lastly used F822 (Figure 1B) and NsgmRNA (1,332 nt) as longer RNA competitors against radiolabeled S2hp. They compete effectively against S2hp-NCAP binding but with some differences (Figure 5G and H). F822 does not disrupt the main S2hp-NCAP complex at 50 nM, reduce it to the complex with two NCAP dimers bound at 100 nM, return some S2hp to the free state at 200 nM, and leave almost all S2hp free at 400 nM (Figure 5G, lanes 3-6). Despite being longer, NsgmRNA needs nearly twice the concentrations in the competition—little disruption at 100 nM, smeary intermediates at 200 nM, partial free S2hp at 400 nM, and almost all free S2hp at 800 nM (Figure 5H, lanes 6–9). These results, as summarized in Figure 5I, are consistent with the fact that F822 is part of the gRNA, which has to outcompete the highly abundant NsgmRNA for packaging during viral replication.

### All three major NCAP regions bind RNA

We expressed and purified a series of truncated NCAP proteins (Figure 1A, Supplementary Table S1) and characterized their RNA-binding properties. N-RBD contains the N-terminal domain to the RNA-binding domain. The central LCD is intrinsically disordered but is soluble. DD-C contains the dimerization domain to the C-terminus. We ran an EMSA of radiolabeled S2hp with NCAP, N-RBD, LCD, and DD-C in the absence and presence of tRNAs (Figure 6A). Without tRNAs, all three NCAP truncations caused the free RNA band to disappear (Figure 6A, lanes 5, 7, 9). N-RBD generated several smeary bands, whereas the complexes with LCD and DD-C were stuck in the wells akin to the S2hp-NCAP complex. The presence of tRNAs returned N-RBD-bound S2hp to the free state (lane 6), indicative of weak interactions. Addition of tRNAs assisted the complexes with LCD and DD-C to migrate into gels (lanes 8, 10). Thus, all three NCAP regions bind viral RNA. Similar EMSAs were performed using radiolabeled S2 (Figure 6B). Unlike NCAP, no truncations were able to bind S2 regardless of tRNAs. This result indicates that these regions work together to achieve high-affinity RNA binding.

**Figure 6.**
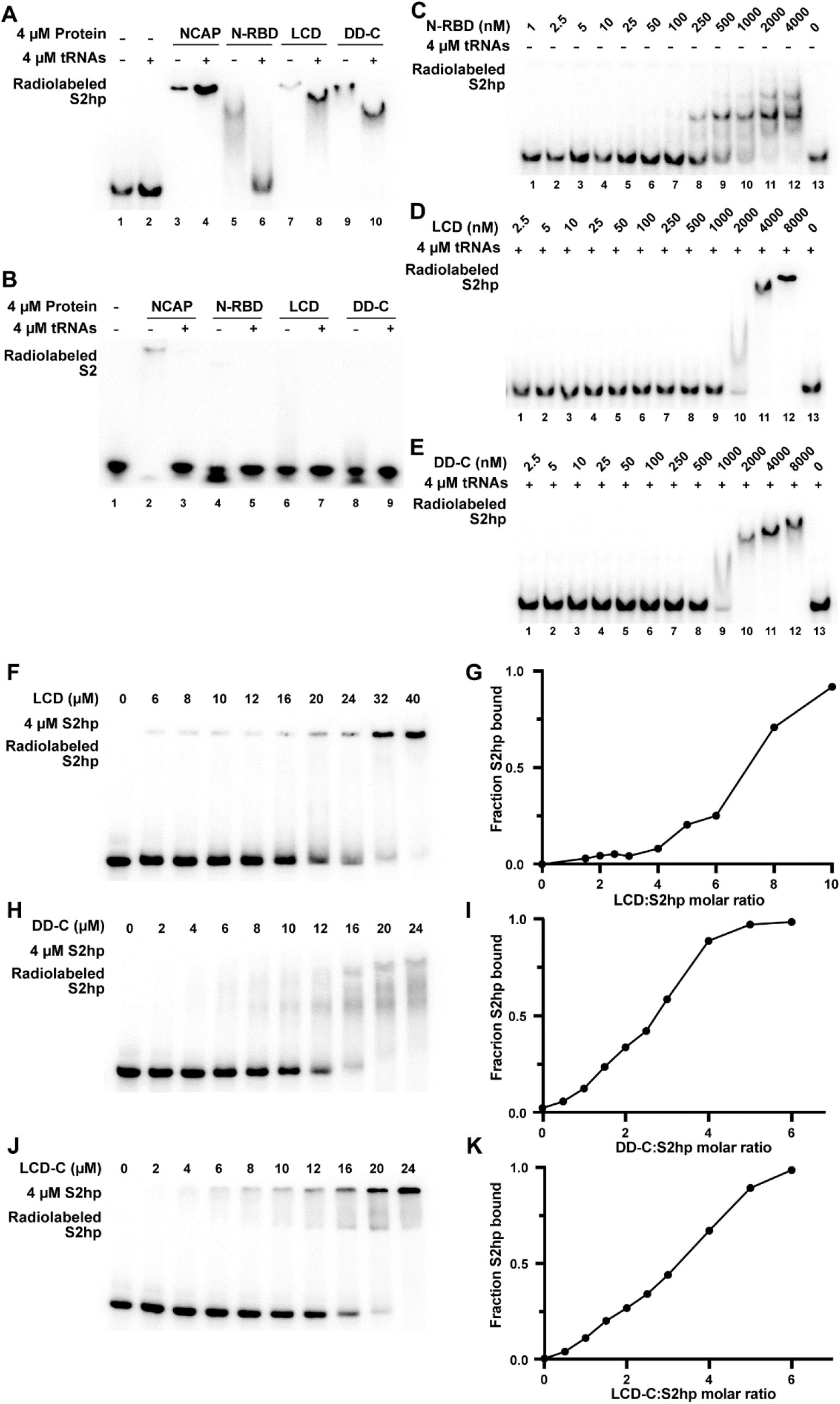
EMSA and protein saturation assays with truncated NCAP constructs N-RBD, LCD, and DD-C. (**A**) NCAP, LCD, and DD-C bind S2hp in the presence and absence of tRNAs while N-RBD only binds S2hp in the absence of tRNA competitors. (**B**) NCAP binds S2 only in the absence of tRNAs. N-RBD, LCD, and DD-C exhibit little to no binding to S2 whether in the presence or absence of tRNAs. (**C**) Gel shift of radiolabeled S2hp with varying concentration of N-RBD and no tRNAs added. (**D**, **E**) Gel shift of radiolabeled S2hp with varying concentration of LCD (**D**) or DD-C (**E**), with 4 μM tRNAs. (**F**) LCD saturation with radiolabeled S2hp, 4 μM unlabeled S2hp, and varying concentrations of LCD. (**G**) Quantification of the S2hp-LCD complex over changing LCD:S2hp ratio. (**H**) DD-C saturation with radiolabeled S2hp, 4 μM unlabeled S2hp, and varying concentrations of DD-C. (**I**) Quantification of the appearance of oligomeric complex with S2hp. (**J**) DD-C saturation with radiolabeled S2hp, 4 μM unlabeled S2hp, and varying concentrations of DD-C shows the appearance of smeary bands, likely due to multiple oligomeric states. (**K**) Quantification of S2hp-DD-C complex.

### Differential contribution of N-RBD, LCD, and DD-C to RNA binding

We estimated the *K*_D_’s of N-RBD, LCD, and DD-C for S2hp. The experiment with N-RBD in the absence of tRNAs showed four complex bands, with the two fastest-migrating bands forming at ~250 nM N-RBD and the slower complexes observed at higher concentrations (Figure 6C). This result indicates that multiple N-RBD proteins can simultaneously bind S2hp, with the estimated *K*_D_ of 250–500 nM for the lower order complexes. The assays for LCD and DD-C were performed with 4 μM tRNAs, which assist the complexes to migrate into gels but decrease the apparent *K*_D_ (Figure 6D). At 2 μM LCD, majority of S2hp was shifted to a smeary band. At 4 and 8 μM, S2hp migrated as single bands with decreasing gel mobility, suggesting that proteins are continuously added to the complex. The *K*_D_ is estimated to be 1–2 μM. Similarly, DD-C shifted S2hp to a smeary band at 1 μM and formed concrete bands at 2, 4, and 8 μM (Figure 6E). These *K*_D_ of DD-C is between 0.5 and 1 μM. Overall, none of three individual regions can completely recapitulate the RNA binding properties of NCAP, with DD-C displaying the strongest S2hp binding.

We measured the S2hp-binding stoichiometry of NCAP truncations. Varying concentrations of truncated proteins were incubated with radiolabeled S2hp and 4 μM S2hp. The results with LCD showed that the LCD-S2hp complex barely migrated into the gel and that approximately 10:1 LCD:S2hp ratio is required to shift all the free RNAs to the complex (Figure 6F, G). This ratio is greater than the 6:1 for NCAP. The gel with DD-C displayed the disappearance of radiolabeled S2hp when the DD-C:S2hp ratio reached 5:1 (Figure 6H, I), which is similar to that of NCAP:S2hp stoichiometry. However, at least three smeary complex bands were observed with no single complex band observed at any DD-C concentration. Finally, we engineered a NCAP construct (called LCD-C) that contains both the LCD and DD-C regions. In the stoichiometry EMSA, LCD-C produced mostly a single band, completely occupying S2hp at 6:1 LCD-C:S2hp ratio (Figure 6J, K). Thus, although LCD and DD-C individually retained some RNA-binding properties of NCAP, both regions are required to assemble a higher-order S2hp-binding complex with proper stoichiometry. We note that LCD-C displayed greater cooperativity than NCAP, as limited intermediate complexes (with one or two dimers bound) were observed. This result suggests that N-RBD reduces RNA-binding cooperativity of NCAP.

### Further increasing viral RNA lengths enhances NCAP-binding affinity

The in-depth analyses of short SARS-CoV-2 fragments demonstrated the preference of NCAP for long RNAs that alternate between structured and single-stranded regions. To test if this preference applies to longer RNAs, we extended S2hp on either or both sides (Figure 1B) and measured the *K*_D_ for NCAP using EMSA (Supplementary Figure S3). F294 (fragment of 294 nt) and F323 are S2hp with the 5’ 83-nt and 3’ 112-nt extensions, respectively, which increased the affinities for NCAP 7- and 5-fold. When both extensions were included, the affinity of the resulting F406 construct was increased 144-fold of that of S2hp, with a *K*_D_ of 1.2 ± 0.2 nM (Table 1). The correlation between affinity and length strongly supports the preference of NCAP for longer viral RNAs. The fact that F294 and F323 cover the whole F406 region and yet none of them can achieve the same affinity rules out the possibility that an uncharacterized site in the extensions is responsible for the 1.2-nM binding.

We further investigated the contribution of the 5’ and 3’ extensions to NCAP binding by fusing them to the neighboring SL4 and SL5, respectively. The 5’ extension-SL4 construct, F127, has an affinity (*K*_D_ of 183 ± 56 nM) roughly doubled that of SL4 alone but only 1/7 that of F294, the 5’ extended S2hp (Table 1). In contrast, the 3’ extension-SL5 construct, F257, has an affinity 6.5 times that of SL5 alone, similar to that of F323, the 3’-extended S2hp. The greater contribution of the 3’ extension may be explained by its larger structures (SL6 and SL7). We note that S2hp, F257, F294, F323, and F406 all contain SL5 and the order of their affinities for NCAP correlates with the number and size of additional structures that can serve as protein-binding sites.

## DISCUSSION

We report several previously unknown RNA-binding properties of the SARS-CoV-2 NCAP. By characterizing various viral and non-viral RNA fragments as models, we discovered that NCAP primarily binds structured RNAs that flank ss NCAP-crosslinking sites. Moreover, NCAP prefers to bind larger structures in the viral genomic RNA, such as the SL5 four-way junction (34), over smaller structures, such as the SL4 hairpin, and over certain non-specific RNAs like tRNAs and P4-P6 even though they are structured. Although it is not known what features of SL5 are important for binding NCAP, it stably folds as a four-way junction that is composed of a long basal stem and three apical stem loops 5a, 5b and 5c (Figure 1B) (34). This structure is phylogenetically conserved among alpha- and beta-coronaviruese (with some exceptions) and has been proposed to function as a packaging signal (by analogy with the transmissible gastroenteritis coronavirus (11)), in regulation of translation (by containing the ORF1a start codon), and possibly as an internal ribosome entry site for directly recruiting ribosome for translation initiation.

Our analyses of NCAP-RNA interaction (Figure 5A–D, and G) provide detailed experimental support to a previous proposal that NCAP prefers multiple structures separated by ss linkers (27). The identification of the flanking structures of S2hp as the primary NCAP-binding sites explains why these structures are important for RNA-binding selectivity of NCAP. NCAP binds viral RNA fragments (S2hp, SL5, SL4, and S2) cooperatively (Figure 2 and Supplementary Figure S1). We think that the cooperativity is achieved by NCAP-NCAP interactions at two levels. In the first level, different NCAP dimers interact with each other upon binding SL5 and S2hp. In the second, any NCAP monomers may dimerize upon binding RNA. RNA-free NCAP has a dimerization constant of 0.4-0.5 μM (21). Binding to the same RNA increases the effective concentration of NCAP monomers to each other. In the presence of tRNAs, the affinity of NCAP for S2hp is higher than those for SL5 and SL4 individually (Figure 2 and Table 1). We suggest that NCAP proteins bound to these structures interact with each other to achieve greater affinity and specificity. The single-stranded linkers between the structures, such as S2, are likely to provide conformational flexibility that allows NCAP proteins bound to neighboring structures to interact with each other. Our competition results showing NCAP binding to structured RNAs suppresses its strong association with ssRNAs does not exclude weak interactions. Quite to the contrary, ssRNA linkers are very likely to interact with NCAP weakly, as detected in the crosslinking study (27) but is unstable under our EMSA conditions.

Our analyses of NCAP domain contribution to RNA binding are consistent with previous reports. We and others have found using non-radiolabeled RNAs that NCAP proteins with one or more major regions deleted have limited to no detectable RNA-binding activities (31, 41). However, Huang and colleagues previously reported that the NTT, LCD, and CTT regions of the SARS-CoV NCAP protein are all capable of binding RNA (35). With radiolabeled RNA, we show RNA-binding activity for all three major NCAP regions and thereby resolved the discrepancy. The N-RBD forms multiple complexes with S2hp that are disrupted by tRNAs (Figure 6C and A), suggesting that this interaction is not very specific. This is somewhat surprising because the RBD of NCAP from the Middle East respiratory syndrome coronavirus (MERS-CoV) can bind a hairpin in the packaging signal (42). Since the packaging signal of SARS-CoV-2 is still elusive, the RNA-binding property of its RBD may be different. The LCD alone can bind viral RNA to form concrete complexes and is required for proper assembly of higher order structures (Figure 6). The interaction between LCD and S2hp withstands competition by tRNAs, demonstrating its surprising strength. Thus, in addition to connect two structured RNA-binding domains and coordinate their interactions with RNA, the LCD plays a greater role in modulating RNA through direct binding. The LCD contains a RS-rich motif in its N-terminal region and several Lys residues neighboring the DD (Figure 1A). These motifs are known to mediate RNA-binding affinity and specificity of other IDRs (43).

Taking these results and previous studies together, we propose a mechanism uniting gRNA packaging and recognition, which we call “recognition-by-packaging.” This mechanism involves two key components. First, NCAP preferentially binds structured RNA elements (Figure 7). The presence of these structures throughout the viral genome are extensively supported by RNA secondary structure prediction, chemical mapping, and electron microscopic studies (44–48). Second, NCAP proteins bound to the same gRNA oligomerize and assemble into higher order structures. Single-stranded regions spaced between NCAP-bound structures provide the conformational flexibility necessary for NCAP proteins to interact with each other favorably in such assemblies. These interactions likely include the previously characterized oligomerization (20, 21, 24, 25) and amyloid-like interactions by the LCD and other IDRs (31, 49). These interactions lead to further stabilization of the NCAP-RNA complex, effectively increasing the binding affinity for the longer gRNAs with optimal structural patterns that can recruit larger number of NCAP proteins. This is a key feature of recognition-by-packaging. Unlike the general thought that NCAP recognizes gRNA and then package it, the central idea of our proposal is that the protein-protein interactions involved in packaging are also critical for RNA recognition. Besides gRNA recognition, our results also suggest that large structures like SL5 and local sites with the favorable structure-ss-structure patterns may be the NCAP-binding sites for its regulatory functions (Figure 7).

**Figure 7.**
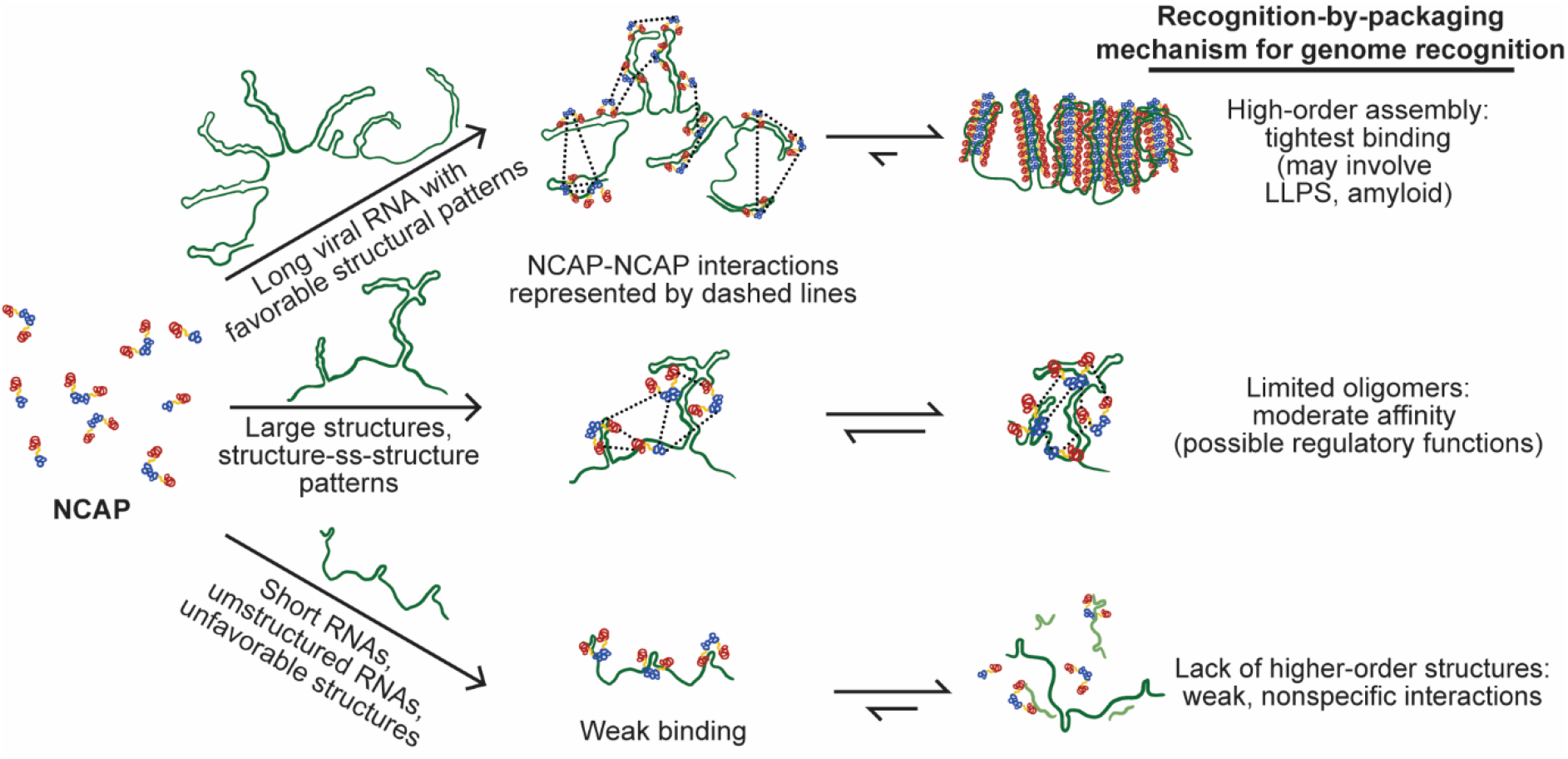
Schematic of how NCAP selects RNA binding partners via oligomerization and higher-order assembly upon association with RNA.

Our proposed mechanism is consistent with recently reported evidence. We show in a parallel work that the NCAP LCD can form amyloid-like fibrils, that RNA greatly stimulates the fibril formation, and that blocking the fibril elongation inhibits viral replication (31). Therefore, these NCAP-NCAP interactions are important for viral functions. The Doudna and Ott groups demonstrated that the 5’ two-thirds of the SARS-CoV-2 genome broadly supports packaging of reporter mRNAs into SARS-CoV-2 virus-like particles produced by co-expression of NCAP and other viral structural proteins (50). This region contains the S2hp sequence and the whole ORF1ab region that are not present in subgenomic mRNAs. This result suggests that selection of viral gRNA may not rely entirely on one contiguous cis-acting packaging signal and is likely to involve the recognition-by-packing mechanism that relies on a network of NCAP-RNA and NCAP-NCAP interactions. On the other hand, a genomic fragment containing nucleotide 20,080–21,171 did display the strongest packaging strength. Although it is unknown whether this fragment is required for viral gRNA packaging, it is likely that the recognition-by-packaging mechanism works in conjunction with high-affinity packaging signals to achieve optimal specificity for the gRNA. We also acknowledge that our study does not include the membrane protein, which could be important for recognizing SARS-CoV-2 gRNA. Further investigation is required to test these possibilities.

Our study lends insight into the LLPS formed by NCAP and other RNA-binding proteins upon binding RNA. LLPS is mediated by multivalent protein-protein and protein-RNA interactions (51, 52). Previous studies suggested that phase separation is driven by specific NCAP-binding elements in the gRNA and that lengthening the RNA by addition of non-viral sequences enhance NCAP LLPS (27). Our biochemical results suggest that a preferred NCAP-binding element is either a single large structure or a cluster of multiple structures separated by ss linkers. Indeed, in a parallel study, we found that compared to S2hp, a higher concentration of S2 is required to induce LLPS with NCAP and its central LCD-containing fragments (31). Recently, Yildiz and colleagues reported that NCAP forms spherical condensates with unstructured RNAs but asymmetric ones with structured viral gRNAs (49). The strong influence of RNA structures on LLPS is further elaborated in a preprint by Roden *et al.,* reporting that dsRNAs (including SL4 and SL5) trigger NCAP to undergo LLPS, whereas ssRNAs are not required (32). These observations may be explained by our findings that structured RNAs bind NCAP better than ssRNAs and that optimal spacing of structures enhances binding to NCAP. Similarly, the detailed understanding of NCAP-RNA interaction explains how gRNAs are selectively recruited to the droplets in the presence of other RNAs and should help investigate the function of LLPS in the viral life cycle.

The RNA recognition mechanism we propose is likely to contribute to other biological processes. Stress granules, P bodies, and nuclear speckles are cellular membraneless organelles made of ribonucleoprotein condensates (51–53). Parker and colleagues reported that larger RNAs accumulate in stress granules (54). No obvious RNA sequences were found to explain the localization. They proposed that long RNAs are enriched in stress granules (and probably other LLPS) due to their ability to forge intermolecular RNA-RNA interactions (55). Our proposed mechanism offers a parallel/alternative explanation. Longer RNAs provide larger numbers of protein-binding sites and thus promote protein-protein interactions that in turn enhance the affinity for these RNAs. A common feature of NCAP and cellular RNA-binding proteins that form membraneless organelles is the presence of functionally important LCDs or IDRs. To fully understand relevant biological functions and diseases, it is essential to integrate the studies of protein-RNA and LCDLCD interactions (56).

Supplementary Data are available.

## ACKNOWLEDGEMENTS

We thank Abby Thurn and Prof. William Gelbart for sharing the Nodamura RNA. We thank Grant Shoffner and Professor Gelbart for comments and discussion.

## FUNDING

This work is supported by the National Science Foundation under Grant No. (MCB 1616265), NIH/NIA R01 Grant AG048120, the U.S. Department of Energy (DOE) Contract No. DOE - DE-FC02-02ER63421, and by UCLA David Geffen School of Medicine – Eli and Edythe Broad Center of Regenerative Medicine and Stem Cell Research Award Program, Broad Stem Cell Research Center (BSCRC) COVID 19 Research Award (OCRC #20-73). E.T-F is supported by the Human Frontiers Science Project Organization (HFSPO) (LT000623/ 2018-L). L.S. is supported by NIH NIGMS GM123126 grant. C.T.Z. is funded by the UCLA Dissertation Year Fellowship. We thank NIH AG AG048120 and HHMI for support.

## Conflict of interest statement

None declared.

## Notes

### Competing Interest Statement

The authors have declared no competing interest.

