## Supplementary data for "The SARS-CoV-2 nucleocapsid protein preferentially binds long and structured RNAs"

### For

**Sequences of protein constructs**

**Sequences of RNAs**

**Supplementary Figures and Tables:**

**Figure S1.** Binding affinities of NCAP for S2hp fragments in the absence of tRNAs.

**Figure S2.** Competition of viral and non-viral RNAs against radiolabeled S2-NCAP binding.

**Figure S3.** Binding affinities of NCAP for longer viral RNA fragments without tRNAs.

**Table S1.** DNA primers used in this work

**Table S2.** Lengths and molecular masses of SARS-CoV-2 and non-SARS-CoV-2 RNAs used.

### ***Sequences of protein constructs (N- to C-termini)***

NCAP

SDNGPQNQRNAPRITFGGPSDSTGSNQNGERSGARSKQRRPQGLPNNTASWFTALTQHGKEDLK
FPRGQGVPINTNSSPDDQIGYYRRATRRIRGGDGKMKDLSPRWYFYYLGTGPEAGLPYGANKDG
IIWVATEGALNTPKDHIGTRNPANNAAIVLQLPQGTTLPKGFYAEGSRGGSQASSRSSSRSRNS
SRNSTPGSSRGTSPARMAGNGGDAALALLLLDRLNQLESKMSGKGQQQQGQTVTKKSAAEASKK
PRQKRTATKAYNVTQAFGRRGPEQTQGNFGDQELIRQGTDYKHWPQIAQFAPSASAFFGMSRIG
MEVTPSGTWLTYTGAIKLDDKDPNFKDQVILLNKHIDAYKTFPPTEPKKDKKKKADETQALPQR
QKKQQTVTLLPAADLDDFSKQLQQSMSSADSTQA

N-RBD

SDNGPQNQRNAPRITFGGPSDSTGSNQNGERSGARSKQRRPQGLPNNTASWFTALTQHGKEDLK
FPRGQGVPINTNSSPDDQIGYYRRATRRIRGGDGKMKDLSPRWYFYYLGTGPEAGLPYGANKDG
IIWVATEGALNTPKDHIGTRNPANNAAIVLQLPQGTTLPKGFYA

LCD

FYAEGSRGGSQASSRSSSRSRNSSRNSTPGSSRGTSPARMAGNGGDAALALLLLDRLNQLESKM
SGKGQQQQGQTVTKKSAAEASKKPRQKR

LCD-C

FYAEGSRGGSQASSRSSSRSRNSSRNSTPGSSRGTSPARMAGNGGDAALALLLLDRLNQLESKM
SGKGQQQQGQTVTKKSAAEASKKPRQKRTATKAYNVTQAFGRRGPEQTQGNFGDQELIRQGTDY
KHWPQIAQFAPSASAFFGMSRIGMEVTPSGTWLTYTGAIKLDDKDPNFKDQVILLNKHIDAYKT
FPPTEPKKDKKKKADETQALPQRQKKQQTVTLLPAADLDDFSKQLQQSMSSADSTQA

DD-C

KPRQKRTATKAYNVTQAFGRRGPEQTQGNFGDQELIRQGTDYKHWPQIAQFAPSASAFFGMSRI
GMEVTPSGTWLTYTGAIKLDDKDPNFKDQVILLNKHIDAYKTFPPTEPKKDKKKKADETQALPQ
RQKKQQTVTLLPAADLDDFSKQLQQSMSSADSTQA

### ***Sequences of RNAs (5' to 3' unless indicated otherwise)***

Lengths and molecular masses are summarized in Table S2.Non-viral residues added to enhance transcription are in lower case.

NsgmRNA

GGUUUAUACCUUCCCAGGUAACAAACCAACCAACUUUCGAUCUCUUGUAGAUCUGUUCUCUAAA
CGAACAAACUAAAAUGUCUGAUAAUGGACCCCAAAAUCAGCGAAAUGCACCCCGCAUUACGUUU
GGUGGACCCUCAGAUUCAACUGGCAGUAACCAGAAUGGAGAACGCAGUGGGGCGCGAUCAAAAC
AACGUCGGCCCCAAGGUUUACCCAAUAAUACUGCGUCUUGGUUCACCGCUCUCACUCAACAUGG
CAAGGAAGACCUUAAAUUCCCUCGAGGACAAGGCGUUCCAAUUAACACCAAUAGCAGUCCAGAU
GACCAAAUUGGCUACUACCGAAGAGCUACCAGACGAAUUCGUGGUGGUGACGGUAAAAUGAAAG
AUCUCAGUCCAAGAUGGUAUUUCUACUACCUAGGAACUGGGCCAGAAGCUGGACUUCCCUAUGG
UGCUAACAAAGACGGCAUCAUAUGGGUUGCAACUGAGGGAGCCUUGAAUACACCAAAAGAUCAC
AUUGGCACCCGCAAUCCUGCUAACAAUGCUGCAAUCGUGCUACAACUUCCUCAAGGAACAACAU
UGCCAAAAGGCUUCUACGCAGAAGGGAGCAGAGGCGGCAGUCAAGCCUCUUCUCGUUCCUCAUC
ACGUAGUCGCAACAGUUCAAGAAAUUCAACUCCAGGCAGCAGUAGGGGAACUUCUCCUGCUAGA

AUGGCUGGCAAUGGCGGUGAUGCUGCUCUUGCUUUGCUGCUGCUUGACAGAUUGAACCAGCUUG
AGAGCAAAAUGUCUGGUAAAGGCCAACAACAACAAGGCCAAACUGUCACUAAGAAAUCUGCUGC
UGAGGCUUCUAAGAAGCCUCGGCAAAAACGUACUGCCACUAAAGCAUACAAUGUAACACAAGCU
UUCGGCAGACGUGGUCCAGAACAAACCCAAGGAAAUUUUGGGGACCAGGAACUAAUCAGACAAG
GAACUGAUUACAAACAUUGGCCGCAAAUUGCACAAUUUGCCCCCAGCGCUUCAGCGUUCUUCGG
AAUGUCGCGCAUUGGCAUGGAAGUCACACCUUCGGGAACGUGGUUGACCUACACAGGUGCCAUC
AAAUUGGAUGACAAAGAUCCAAAUUUCAAAGAUCAAGUCAUUUUGCUGAAUAAGCAUAUUGACG
CAUACAAAACAUUCCCACCAACAGAGCCUAAAAAGGACAAAAAGAAGAAGGCUGAUGAAACUCA
AGCCUUACCGCAGAGACAGAAGAAACAGCAAACUGUGACUCUUCUUCCUGCUGCAGAUUUGGAU
GAUUUCUCCAAACAAUUGCAACAAUCCAUGAGCAGUGCUGACUCAACUCAGG

F822

ggAUUAAAGGUUUAUACCUUCCCAGGUAACAAACCAACCAACUUUCGAUCUCUUGUAGAUCUGU
UCUCUAAACGAACUUUAAAAUCUGUGUGGCUGUCACUCGGCUGCAUGCUUAGUGCACUCACGCA
GUAUAAUUAAUAACUAAUUACUGUCGUUGACAGGACACGAGUAACUCGUCUAUCUUCUGCAGGC
UGCUUACGGUUUCGUCCGUGUUGCAGCCGAUCAUCAGCACAUCUAGGUUUCGUCCGGGUGUGAC
CGAAAGGUAAGAUGGAGAGCCUUGUCCCUGGUUUCAACGAGAAAACACACGUCCAACUCAGUUU
GCCUGUUUUACAGGUUCGCGACGUGCUCGUACGUGGCUUUGGAGACUCCGUGGAGGAGGUCUUA
UCAGAGGCACGUCAACAUCUUAAAGAUGGCACUUGUGGCUUAGUAGAAGUUGAAAAAGGCGUUU
UGCCUCAACUUGAACAGCCCUAUGUGUUCAUCAAACGUUCGGAUGCUCGAACUGCACCUCAUGG
UCAUGUUAUGGUUGAGCUGGUAGCAGAACUCGAAGGCAUUCAGUACGGUCGUAGUGGUGAGACA
CUUGGUGUCCUUGUCCCUCAUGUGGGCGAAAUACCAGUGGCUUACCGCAAGGUUCUUCUUCGUA
AGAACGGUAAUAAAGGAGCUGGUGGCCAUAGUUACGGCGCCGAUCUAAAGUCAUUUGACUUAGG
CGACGAGCUUGGCACUGAUCCUUAUGAAGAUUUUCAAGAAAACUGGAACACUAAACAUAGCAGU
GGUGUUACCCGUGAACUCAUGCGUGAGCUUAACGGAGGGGCAUACACUCGCUAUGU

F406

ggAUUAAAGGUUUAUACCUUCCCAGGUAACAAACCAACCAACUUUCGAUCUCUUGUAGAUCUGU
UCUCUAAACGAACUUUAAAAUCUGUGUGGCUGUCACUCGGCUGCAUGCUUAGUGCACUCACGCA
GUAUAAUUAAUAACUAAUUACUGUCGUUGACAGGACACGAGUAACUCGUCUAUCUUCUGCAGGC
UGCUUACGGUUUCGUCCGUGUUGCAGCCGAUCAUCAGCACAUCUAGGUUUCGUCCGGGUGUGAC
CGAAAGGUAAGAUGGAGAGCCUUGUCCCUGGUUUCAACGAGAAAACACACGUCCAACUCAGUUU
GCCUGUUUUACAGGUUCGCGACGUGCUCGUACGUGGCUUUGGAGACUCCGUGGAGGAGGUCUUA
UCAGAGGCACGUCAACAUCUUAAA

F323

ggCUGUGUGGCUGUCACUCGGCUGCAUGCUUAGUGCACUCACGCAGUAUAAUUAAUAACUAAUU
ACUGUCGUUGACAGGACACGAGUAACUCGUCUAUCUUCUGCAGGCUGCUUACGGUUUCGUCCGU
GUUGCAGCCGAUCAUCAGCACAUCUAGGUUUCGUCCGGGUGUGACCGAAAGGUAAGAUGGAGAG
CCUUGUCCCUGGUUUCAACGAGAAAACACACGUCCAACUCAGUUUGCCUGUUUUACAGGUUCGC
GACGUGCUCGUACGUGGCUUUGGAGACUCCGUGGAGGAGGUCUUAUCAGAGGCACGUCAACAUC
UUAAA

F294

ggAUUAAAGGUUUAUACCUUCCCAGGUAACAAACCAACCAACUUUCGAUCUCUUGUAGAUCUGU
UCUCUAAACGAACUUUAAAAUCUGUGUGGCUGUCACUCGGCUGCAUGCUUAGUGCACUCACGCA
GUAUAAUUAAUAACUAAUUACUGUCGUUGACAGGACACGAGUAACUCGUCUAUCUUCUGCAGGC

UGCUUACGGUUUCGUCCGUGUUGCAGCCGAUCAUCAGCACAUCUAGGUUUCGUCCGGGUGUGAC
CGAAAGGUAAGAUGGAGAGCCUUGUCCCUGGUUUCAACGA

F257
ggUCGUUGACAGGACACGAGUAACUCGUCUAUCUUCUGCAGGCUGCUUACGGUUUCGUCCGUGU
UGCAGCCGAUCAUCAGCACAUCUAGGUUUCGUCCGGGUGUGACCGAAAGGUAAGAUGGAGAGCC
UUGUCCCUGGUUUCAACGAGAAAACACACGUCCAACUCAGUUUGCCUGUUUUACAGGUUCGCGA
CGUGCUCGUACGUGGCUUUGGAGACUCCGUGGAGGAGGUCUUAUCAGAGGCACGUCAACAUCUU
AAA

F127
ggAUUAAAGGUUUAUACCUUCCCAGGUAACAAACCAACCAACUUUCGAUCUCUUGUAGAUCUGU
UCUCUAAACGAACUUUAAAAUCUGUGUGGCUGUCACUCGGCUGCAUGCUUAGUGCACUCACGCA
G

S2hp
GGCUGUGUGGCUGUCACUCGGCUGCAUGCUUAGUGCACUCACGCAGUAUAAUUAAUAACUAAUU
ACUGUCGUUGACAGGACACGAGUAACUCGUCUAUCUUCUGCAGGCUGCUUACGGUUUCGUCCGU
GUUGCAGCCGAUCAUCAGCACAUCUAGGUUUCGUCCGGGUGUGACCGAAAGGUAAGAUGGAGAG
CCUUGUCCCUGGUUUCAACGA

SL5
GGUCGUUGACAGGACACGAGUAACUCGUCUAUCUUCUGCAGGCUGCUUACGGUUUCGUCCGUGU
UGCAGCCGAUCAUCAGCACAUCUAGGUUUCGUCCGGGUGUGACCGAAAGGUAAGAUGGAGAGCC
UUGUCCCUGGUUUCAACGA

SL4
GGCUGUGUGGCUGUCACUCGGCUGCAUGCUUAGUGCACUCACGCAG

S2
UAUAAUUAAUAACUAAUUACUG

S1hp
GGUUUAUACCUUCCCAGGUAACAAACCAACCAACUUUCGAUCUCUUGUAGAUC

S1.5hp
GGAUCUCUUGUAGAUCUGUUCUCUAAACGAACUUUAAAAUCUGUGUGGCUGUCACUCGGCUGCA
UGCUUAGUGCACUCACGCAG

S8hp
GGAAGGAGCUGGUGGCCAUAGUUACGGCGCCGAUCUAAAGUCAUUUGACUUAGGCGACGAGCUU
GGCACUGAUCCUUAUGAAGAUUUUCAAGAAAACUGGAACACUAAACAUAGCAGUGGUGUUACCC
GUGAACUCAUGCGUGAGCUUAACGGAGGGGCAUACACUCGCUAUGU

S1
AACCAACUUUC

S1.5
UGUUCUCUAAACGAACUUUAAAAU

S8
AUGAAGAUUUUCAAGAAAACUGGAACACUAA

Nodamura replicon
GUAUUGAAUCCAAAACUCAAAAUGCUGAACUACGAGACAAUCAUCAACGGCGCAUCGAGCGCUC
UGAACAUCGUUUCGCGUGCGUUAGGAUACCGCGUGCCACUAGCCAAAUCGCUGGCGCUGGUCGC
GGGGUCCUGCGUGGUGUACAAAAUAAUCGUGCAUCGACGCACGCUCGUGGCGUUCCUGGUAAUC
GGACCAUACGCCACGGUGGUGCAGCACCGUCUGCCGAUGGCCCUUCAGAGGGCCAUCAUUGAAU
AUACACGAGAAGACCGUGAGAUCAGCCUGUUUCCGCAAAAUUCCAUCGUUUCCGCAGAACACGC
GCGGAAAGCGGAUAAUGGGCAUCCGAUCUCCGGGGGAACGCGUGAUGUCGCGAGGGAGACUAUU
UCCCUUGCCAUAAGGGCCGCUGGUUUUCGUCAUUACGAAAUCAGCCCCGCGAGGCAAUCACCAG
CUGAGGCGGCAAGCCACCAACAUUAUGCCGCCGCUGACCUCGUGAGAGCGGCUACUGAAGAUAA
GAUCCAAGACGGUGAUGUGGUAGUUGCCAUCGACAUCGAUUACUACCUGCGUGACAUUGACCGC
UACCUGGGUCGUGGUGUCCCGUUCAUGGCUUACACCUUCAAUCCUGUUGAAGUAGCUGGCCGUG
ACGGUGACUCCUUUUUCCGGAUCACGAACAAUCAAGUCACGUUUGAUGUUAGUGGCGGUGGAUC
UUGGUCCCAUGAAGUUUGGGACUGGUGCGCGUUUGGUGAGUUCAUCGAGACCCGAGACGCGAGC
UGGCUUGCUUGGUUCGCCCGGGCGGUUGGACUCACCAAGUCGCAGAUCCACAAAGUUCACUACU
GCCGUCCAUGGCCGCAAUCGCCCCAUCGCGCUUUGGUGUGGUGUCUGCCUGUAGCAAGCUACUG
GCGCUUCACUUUCAUUCCGACGGACCUGCAUACGCGCACGCUUCGGCGUGUGCGUUAUCAGGAC
ACGUCCCGGCCCGGUUGGAAUUCCAUCGUCUCGACCGGGUCCGAAGGCCUUAAUAUCAGCCUUG
GUCGCGAAGGAGCUGAUCAUUGCGUGACGAUUCCAAAGGUGCACUACGACAUGCUUAUGGGUUU
GUCGAGUGCGCAGUCGUUGUCGUCCCGCAUGAUCGGGCUCAAGUACACUGAUCCUAGUGUACUC
GCGACGGUUGCCCAAUACUAUCAGGGCAAGAAUGUUGAAGUUGCCGACGCUGACAGGAUCGGCC
GCGCCAUAAAUCCCAAGGUCCACUGGCCAGCGCACGUCGAAGUUGACGAGGCGGAGGUUAGUGC
UCGGGUGUACGCCAGCCCGUUGGUAUCUGACGAAAAUAUGAUGCCUAUGAUCAAGCGCUGGGAG
ACGCUGUCGUUGUCGCUGGACCGCCGGGUUACAUUCCAACGUAAUCCGAAGGUUCCUGGAAAAC
GGCUCAGGGCUUAUGCCAUUGAGUUCGUUGACUUGGUUGUGCCUGAGCGUGGUGUCGGAGUCCC
CUAUUCAUUGGAGGACACCGCCGCCAUGCUGGACAAACCAAGCCAGACCCUCGCCAUCCAACAG
GUGUGGGAGACUGUCGACAUGCCCCCAAGAAGGCUCAUCGAAGCGUUCGUGAAGAACGAACCGA
CCAUGAAGGCUGGCCGUAUCAUCUCGUCGUUCGCUGACAUGCGGUUCCUACUGCGGUUUUCCAG
CUAUACGCUGGCAUUCCGUGAUCAGGUGCUGCAUGCAGAGCACAACCGGCAUUGGUUUUGCCCG
GGUUUGACCCCCGAGCAGAUCGCCACAAAAGUGGUUGAUUACGUGUCCGGUGUUGAAGAACCAU
CGGAGGGAGACUUUUCCAACUUUGAUGGCACGGUUAGUGAGUGGCUACAACGCCACGUCAUGAA
CGCCGUCUACCUGCGUUAUUUCAAUCACCGAGCGCAGCGAGACCUCAGGUCGUAUACCGACAUG
CUGGUCUCAUGCCCCGCGAGGGCGAAGCGAUUCGGUUUUGCUUAUGACGCGGGUGUCGGCGUUA
AGAGCGGGUCGCCAACAACUUGCGACCUGAAUACCGUGUGCAAUGGUUUCCUCCAAUAUUGCUC
CAUUCGAAUGACACACCCAGAGCUGACACCAAUCGAUGCUUUCCGGCUCAUCGGUCUCGCGUUU
GGGGACGAUUCCCUCUUCGAGCGACGUUUCGCUAAGAACUAUGCGAAGGUUUCCGCCGAGGUGG
GGAUGGUCCUCAAAAUCGAGCGAUUCGACCCGGCACAAGGCAUCACUUUCCUCGCCCGUGUUUA
UCCCGACCCCUACACGUCGACCACAAGUUUCCAGGACCCUUUGCGUACCUGGAGGAAGCUCCAC
UUGACGACGCGCGAUCCAACAAUACCAUUGGCAACGGCUGCCAUCGAUCGCGUUGAGGGCUACC
UCGUCACCGACGGCCUGAGCCCGCUUACUGGCGCGUAUUGUCGCAUGGUUAAGCGGGUUUACGA
GGCCGGCGGAGCCGAGGAUGCCGCCAAGAGGAGGUCGCGAAAAUCCCAUUCCCGCGAAAAGCCG
UAUUGGUUGACUGUUGGAGGCGCUUGGCCCCAAGAUGUCAAGGACGUUGAUCUUAUGUUCCAGU

GUGCGGCCGCACGUACCGGAGUAGACCUCGAGACACUUCGGUCUCUGGAUCAGCGUCUAGGAGA
AAUCACUGACGUCUGGGCGGAUAUUACCAUCAACCGGGAUAAUGAACCAAACCCCUACAAGGAU
ACACUGGACUUGGAGGGCCCGGCUGAUGGCCGGGUGGACGAUCGUGUAUUUCAGAAUGACAAAC
AUGUCAUGAGGUUAAGAGCUAAUCAAGUCACUUCCAGCCAAGCUGGAGCAGCUGGCUCAGGAGA
CGCAAGCAACGAUCCAAACGCUCAUGAUCGCGGAUCCCAACGUCAACAAGGAUCUGCGAGCGUU
CUGCGAGUUCCUGACCGUGCAGCACCAGCGGGCGUAUCGAGCGACGAACAGCCUGCUCAUCAAA
CCGCGAGUCGCAGCAGCGCUUCGCGGGGAGGAGCUGGACCUGGGCGAGGCGGACGUCGCCGCCC
GGGUCCGCCAGCUAAAACAACAGCUGGCGACGAGAUGGAAAUCAAGCCAGGGCACCAACAAGUG
GCCCGUCAAAACGCCAAGCUGAGGGAAGGUCCAGAUCUAGUCGAGGACCGGCGGGUAGCCGUGG
ACGCGGGAAAGACUAUAAGGAGGGUGGCAGCGGAGCUAGCGGAGUCAGAGCUGAAGGUAGAGGC
UCUUUGCUCACCUGUGGAGAUGUGGAAGAGAACCCUGGACCC

<u>Tet3-9</u>
GGACCGUCAAAUUGCGGGAAAGGGGUCAACAGCCGUUCAGUACCAAGUCUCAGGGGAAACUUUG
AGAUGGCCUUGCAAAGGGUAUGGUAAUAAGCUGACGGACAUGGUCCUAACCACGCAGCCAAGUC
CUAAGUCAACAGGAGACUGUUGAUAUGGAUGCAGUUCACAGACUAAAUGUCGGUCGGGGAAGAU
GUAUUCUUCUCAUAAGAUAUAGUCGGACCUCUCCCGAAAGGGAGUUGGAGUACUCGAAAUUCGU
UUGGAGGGCUGCAGGGCCCUCUAGA

<u>P4-P6</u>
GGAAUUGCGGGAAAGGGGUCAACAGCCGUUCAGUACCAAGUCUCAGGGGAAACUUUGAGAUGGC
CUUGCAAAGGGUAUGGUAAUAAGCUGACGGACAUGGUCCUAACCACGCAGCCAAGUCCUAAGUC
AACAGAUCUUCUGUUGAUAUGGAUGCAGUUCA

<u>siDGCR8-1 sense</u>
p-CAUCGGACAAGAGUGUGAUUU (p presents 5′-phosphate)

<u>siDGCR8-1 antisense</u>
p-AUCACACUCUUGUCCGAUGUU

<u>siDGCR8-1 duplex</u>
 5′p-CAUCGGACAAGAGUGUGAUUU-3′
3′-UUGUAGCCUGUUCUCACACUA-p5′

### Supplementary Figures and Tables

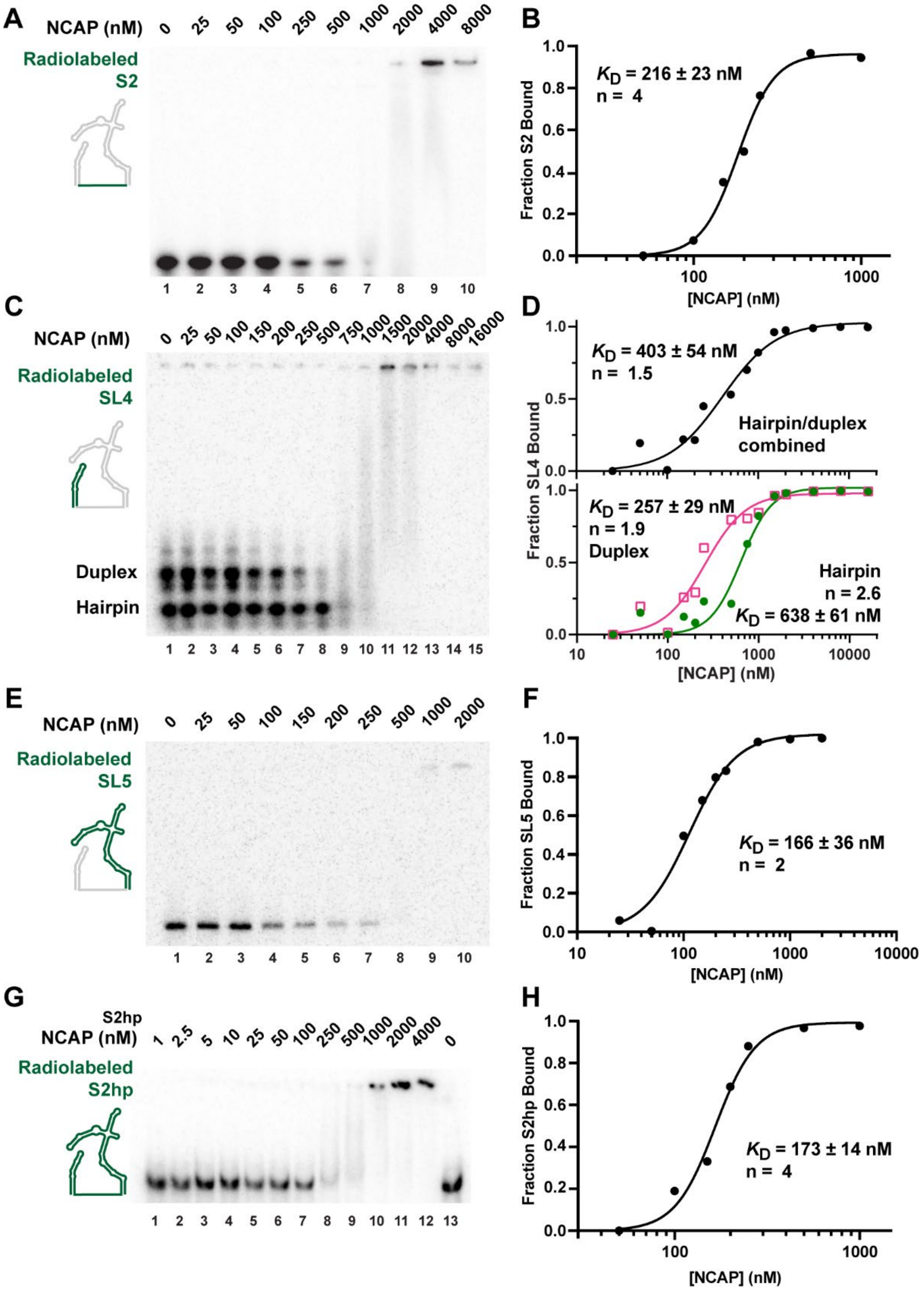


**Figure S1.** Binding affinities of NCAP for S2hp fragments in the absence of tRNAs. The radiolabeled RNA fragments used are highlighted in green in the schematics. Without tRNAs, all complexes with NCAP do not migrate reliably into gels. Binding is inferred from disappearance of the free RNAs. (**A**) EMSA of radiolabeled S2 with varying concentrations of NCAP. Quantification and fitting results are shown in (**B**). (**C**) EMSA of radiolabeled SL4. Quantification and fitting results are shown in (**D**). The free SL4 bands were quantified either combined (top panel) or separately as duplex and hairpin (bottom panel). The Hill coefficient was allowed to vary during the fitting. (**E**) EMSA of radiolabeled SL5. Quantification and fitting results are shown in (**F**). (**G**) EMSA of radiolabeled S2hp. Quantification and fitting results are shown in (**H**). The experiments were repeated 3–7 times and the results are summarized in Table 1. Shown here are representative gel images and quantification results. Curving fitting provides the $K_D$ values with standard errors.

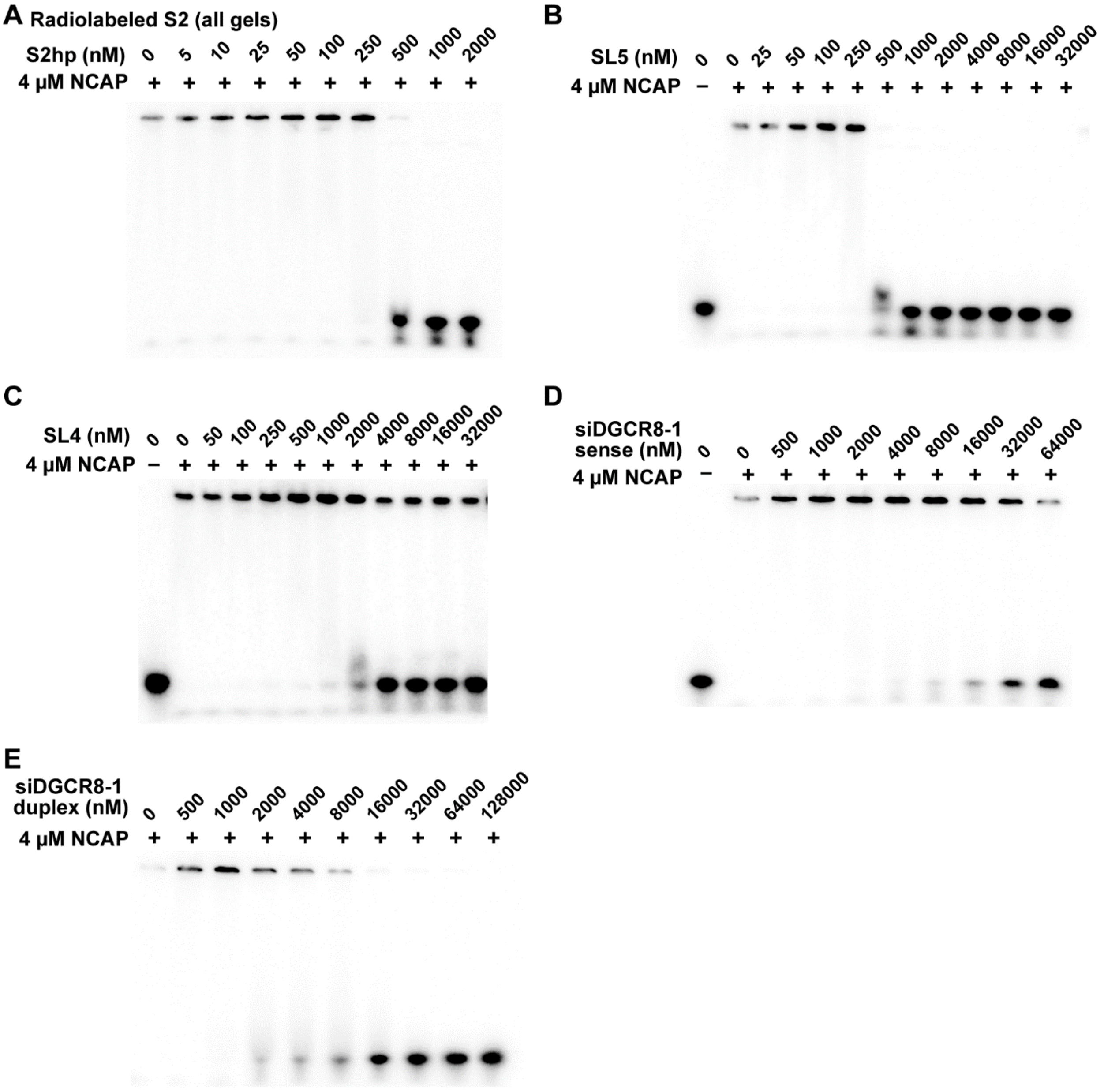


**Figure S2.** Competition of viral and non-viral RNAs against radiolabeled S2-NCAP binding.
Unlabeled S2hp (**A**), SL5 (**B**), SL4 (**C**), siDGCR8-1 sense strand (**D**), and siDGCR8-1 duplex (**E**) were added at the indicated molar concentrations.

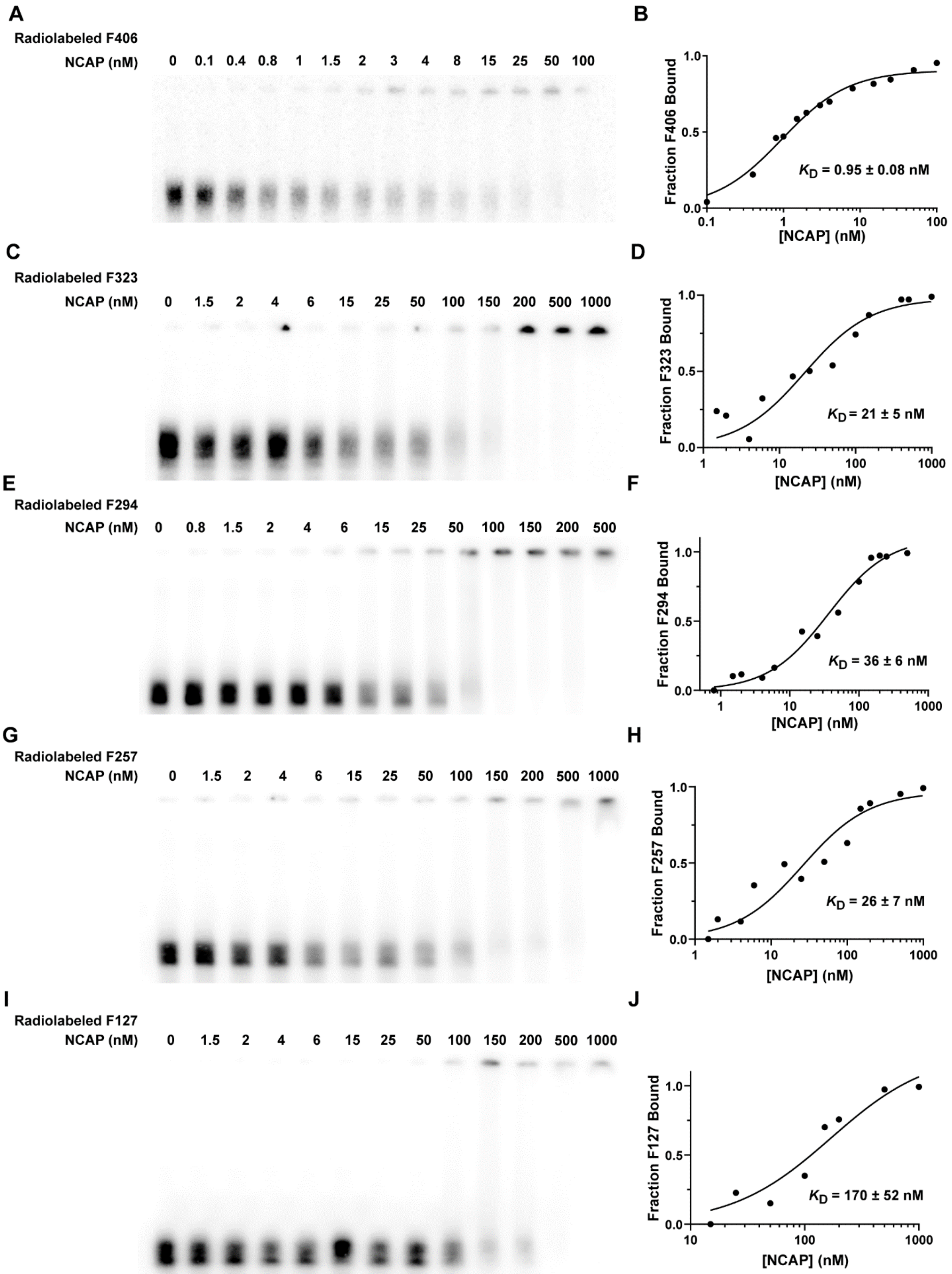


**Figure S3.** Binding affinities of NCAP for longer viral RNA fragments without tRNAs. Similar to Figure S1.

**Table S1.** DNA primers used in this work

| Name* | Primer (5′ to 3′) |
|---|---|
| F822 forward<br>F406 forward<br>F294 forward<br>F127 forward | GCTCTAGA**TAATACGACTCACTATA**GGATTAAAGGTTTATACCTTCCCAG<br>*XbaI* **T7 promoter** |
| 5′-end leader to NsgmRNA | ACAAACTAAAATGTCTGATAATGGACCCCA |
| F416 forward | **TAATACGACTCACTATA**GGATGGCACTTGTGGCTTAGT |
| S1hp forward<br>NsgmRNA forward | **TAATACGACTCACTATA**GGTTTATACCTTCCCAGGTAAC |
| S1.5hp forward | **TAATACGACTCACTATA**GGATCTCTTGTAGATCTGTTCTC |
| S2hp forward<br>SL4 forward<br>F323 forward | **TAATACGACTCACTATA**GGCTGTGTGGCTGTCACTCG |
| SL5 forward | biotin-**TAATACGACTCACTATA**GGTCGTTGACAGGACACGAGT |
| S8hp forward | **TAATACGACTCACTATA**GGAAGGAGCTGGTGGCCATAG |
| NsgmRNA reverse | CATGGTACCCTGAGTTGAGTCAGCACTG<br>KpnI |
| F406 reverse<br>F323 reverse<br>F257 reverse | TTTAAGATGTTGACGTGCCTC |
| F416 reverse | ACATAGCGAGTGTATGCCCC |
| S1hp reverse | TGCACTGCAGATCTACAAGAGATCGAAAG<br>PstI |
| S1hp reverse | GATCTACAAGAGATCGAAAG |
| S1.5hp reverse primer with KpnI | CGGGGTACCTGCGTGAGTGCACTAAG<br>KpnI |
| S1.5hp reverse<br>SL4 reverse<br>F127 reverse | CTGCGTGAGTGCACTAAG |
| S2hp reverse | CGGGGTACCTCGTTGAAACCAGGGACAAG<br>KpnI |
| S2hp reverse | TCGTTGAAACCAGGGACAAG |

| | |
|---|---|
| SL5 reverse<br>F294 reverse | |
| S8hp reverse<br>F822 reverse | ACATAGCGAGTGTATGCCC |
| Biotinylated T7 promoter | biotin-**TAATACGACTCACTATA**GG |

* Some PCR primers were used to generate multiple clones or transcription templates. In such cases, all construct names are listed.

**Table S2.** Lengths and molecular masses of SARS-CoV-2 and non-SARS-CoV-2 RNAs used.

| *RNAs Used* | *Length (nt)* | *Molecular Mass (Da)* |
|---|---|---|
| **SARS-CoV-2 RNAs:** | | |
| NsgmRNA | 1,332 | 428,475 |
| F822* | 824 | 264,826 |
| F406* | 408 | 130,732 |
| F323* | 325 | 104,342 |
| F294* | 296 | 94,714 |
| F257* | 259 | 83,235 |
| F127* | 129 | 41,122 |
| S2hp | 213 | 68,324 |
| SL5 | 147 | 47,217 |
| SL4 | 46 | 14,733 |
| S2 | 22 | 6,941 |
| S1hp | 53 | 16,768 |
| S1.5hp | 84 | 26,793 |
| S8hp | 174 | 56,155 |
| S1 | 11 | 3,394 |
| S1.5 | 24 | 7,567 |
| S8 | 31 | 9,966 |
| **Non-SAR-CoV-2 RNAs:** | | |
| Nodamura replicon | 3,212 | 1,044,882 |
| Tet3-9 | 281 | 90,911 |
| P4-P6 | 160 | 51,708 |
| siDGCR8-1 sense | 21 | 7,037 |
| siDGCR8-1 antisense | 21 | 6,871 |
| siDGCR8-1 duplex | 42 | 13,908 |

* Two non-viral G residues were added at the 5′ ends of these RNAs to facilitate T7 transcription. The RNA fragments were named based on the number of viral residues in their sequences.
